# Processing of a spoken narrative in the human brain is shaped by family cultural background

**DOI:** 10.1101/2020.05.11.083931

**Authors:** M. Hakonen, A. Ikäheimonen, A. Hultèn, J. Kauttonen, M. Koskinen, F-H. Lin, A. Lowe, M. Sams, I. P. Jääskelainen

## Abstract

Using neuroimaging, we studied influence of family cultural background on processing of an audiobook in human brain. The audiobook depicted life of two young Finnish men, one with the Finnish and the other with the Russian family background. Shared family cultural background enhanced similarity of narrative processing in the brain at prelexical, word, sentence, and narrative levels. Similarity was also enhanced in brain areas supporting imagery. The cultural background was further reflected as semantic differences in word lists by which the subjects described what had been on their minds when they heard the audiobook during neuroimaging. Strength of social identity shaped word, sentence, and narrative level processing in the brain. These effects might enhance mutual understanding between persons who share family cultural background and social identity and, conversely, deteriorate between-group mutual understanding in modern multicultural societies wherein native speakers of a language may assume highly similar understanding.

## Introduction

People raised and living in a shared cultural environment and using the same language largely share similar knowledge, experiences, beliefs, values, attitudes and interaction rules that together facilitate shared understanding and fosters smooth cooperation. These similarities create “homophily” – love of the same – that has been studied in sociology for decades using behavioral methods (*1*). In a recent landmark study, Parkinson et al. (*2*) provided neuroimaging evidence for homophily showing that brain activity is exceptionally similar among friends – in contrast to acquaintances – when they are viewing audiovisual film clips. The interest toward the effects of cultural homophily on neural mechanisms of cognitive functions has increased rapidly, as exemplified in the emergence of the new field of “cultural neuroscience” (*3, 4*). Previous behavioral (*5–10*) and neuroimaging (*11*) studies, using pictures as stimuli, have shown that persons living in Eastern (Asian) and Western (American) cultures exhibit robust within-culture similarities and between-culture differences in their perceptual-cognitive styles (*3, 4*). A study that investigated perceptual-cognitive styles of Asian Americans and European Americans found that those of Asian Americans’ were either identical to those of European Americans’ or fell in-between the styles of East Asians’ and European Americans’ (*12*). However, the effects of shared cultural family backgrounds within modern multi-ethnic nations are less well known than those arising in inter-national/continental comparative studies. In multicultural societies native speakers of the same language may assume highly similar understanding of concepts (*13*). It might go unnoticed that this is not true and, therefore, might cause communication mismatches.

Furthermore, since previous cultural neuroscience studies have mostly used pictures as stimuli, it is also less clear whether and how shared family cultural background shapes processing and understanding of natural speech in the brain at different hierarchical levels of language processing. Such processing levels include prelexical processing, which is taking place in the auditory cortices at superior aspects of temporal lobes, and when individual words or short sentences are used as stimuli more pronounced brain activity has been described in left-hemisphere lateral and anterior temporal cortical areas (*14*). In contrast, presentation of a narrative in naturalistic experimental settings results in more widespread and bilateral activation involving prefrontal and parietal areas as well cingulate cortices and precuneus that has been specifically associated with understanding of the story lines in narratives (*15–19*). Visual cortex is also activated during listening to a narrative, likely related to visual imagery elicited by the narrative (*20*).

Previous behavioral cross-cultural studies have provided evidence that cultural background can shape understanding of culturally specific aspects in narratives (*21*). An event related response study by Ellis et al. (*22*) provided related results by showing that the so-called N400 event-related potential response amplitude of fluent Welsh-English bilinguals was significantly stronger to sentences written in Welsh than to sentences written in English. Importantly, this difference was only found for the sentences that contained information about Wales culture but not for the sentences that did not contain culture-related information. The authors interpreted this to reflect that language interacts with factors related to personal identity, such as culture, to shape processing of incoming semantic information. Notably, similarities-differences in language processing might play an important role in mutual understanding and homophily between individuals with shared cultural backgrounds and also in failures of mutual understanding between individuals from different cultural backgrounds.

Recent methodological advances have made it possible to both record brain activity (*2, 15, 17, 18, 23–26*) and behaviorally assess subjective interpretations (*20*) during listening to an audiobook. Here, we thrived on these advances to investigate whether the family cultural background modulates inter-subject similarity in how an audiobook is interpreted, and how the audiobook is processed at multiple different hierarchical levels of language processing in the human brain. We used dynamic inverse functional magnetic resonance imaging (fMRI) (*27*) whose ultra-fast 10-Hz sampling rate we regarded as helpful in studying more accurately the processing of natural speech wherein individual words are presented at rates of about three words per second. We recruited 48 healthy residents of Finland, all fluent in Finnish, to the study. Half of the subjects had Finnish parents and either one or both parents of the other half of the subjects were Russians. The subjects listened to a 71-min audiobook, a prose depicting life of two young Finnish men, one with the Finnish and the other with the Russian family background, during the fMRI. In the audiobook, social interaction scenes were interspersed with descriptions of city scenery amidst changing seasons. Afterwards, the subjects performed an association task on the story replayed in short segments where they were asked to produce words describing what the previous segment had brought to their minds during the neuroimaging session. By mapping the words to semantic vector space (*28*), we estimated the similarity of the meanings and imagery elicited by the story (*20*). Additionally, we measured, using Implicit Association Test (IAT), whether the subject unconsciously associate her/his family’s culture to the more positive attributes than the other culture (*29*). The subjects also filled a questionnaire with items relevant to the family cultural background, including how Russian the subjects self-identify themselves and how well they speak Russian in addition to Finnish. We hypothesized to find family cultural background specific enhancements of similarity in how different parts of the story are interpreted as well as in similarity of brain hemodynamic activity in brain areas known to be involved in processing of natural speech to pinpoint the level(s) of hierarchy of language processing (prelexical *>* word meanings *>* story line) and visual imagery that family cultural background might shape.

## Results

### Cultural family background increased similarity of audiobook interpretation

We estimated similarities in interpretation of the audiobook using the association task. Subjects listened to the audiobook in 101 segments and, after each segment, typed in the following 20–30 sec a list of words that best described what had been on their minds at that point in the audiobook during neuroimaging (Fig. 1; for the description of the method, see also (*20*)). The semantic similarity of the subjects’ word lists was estimated by transforming the lists into the vector representations in a semantic space and by calculating the cosine similarities between the resulting semantic vectors (*28*). There were significant effects both in segments depicting interacting protagonists and in segments with descriptions of city scenery amidst changing seasons. The Russian-background subjects listed 33% more words than the Finnish-background subjects (18 724 *vs*. 12 536 words; p < 0.01, as assessed with a paired T-test). The similarity was significantly higher in 44/101 segments (p<0.05, as assessed with a paired t-test, false discovery rate (FDR) -adjusted). However, in Finnish-background *vs*. Russian-background subjects the similarity was significantly higher in 12 out of 101 segments (p<0.05, FDR-adjusted). The segments where there were significant effects are indicated in the English-translated transcription of the audiobook in Supplementary Materials.

**Fig. 1.**
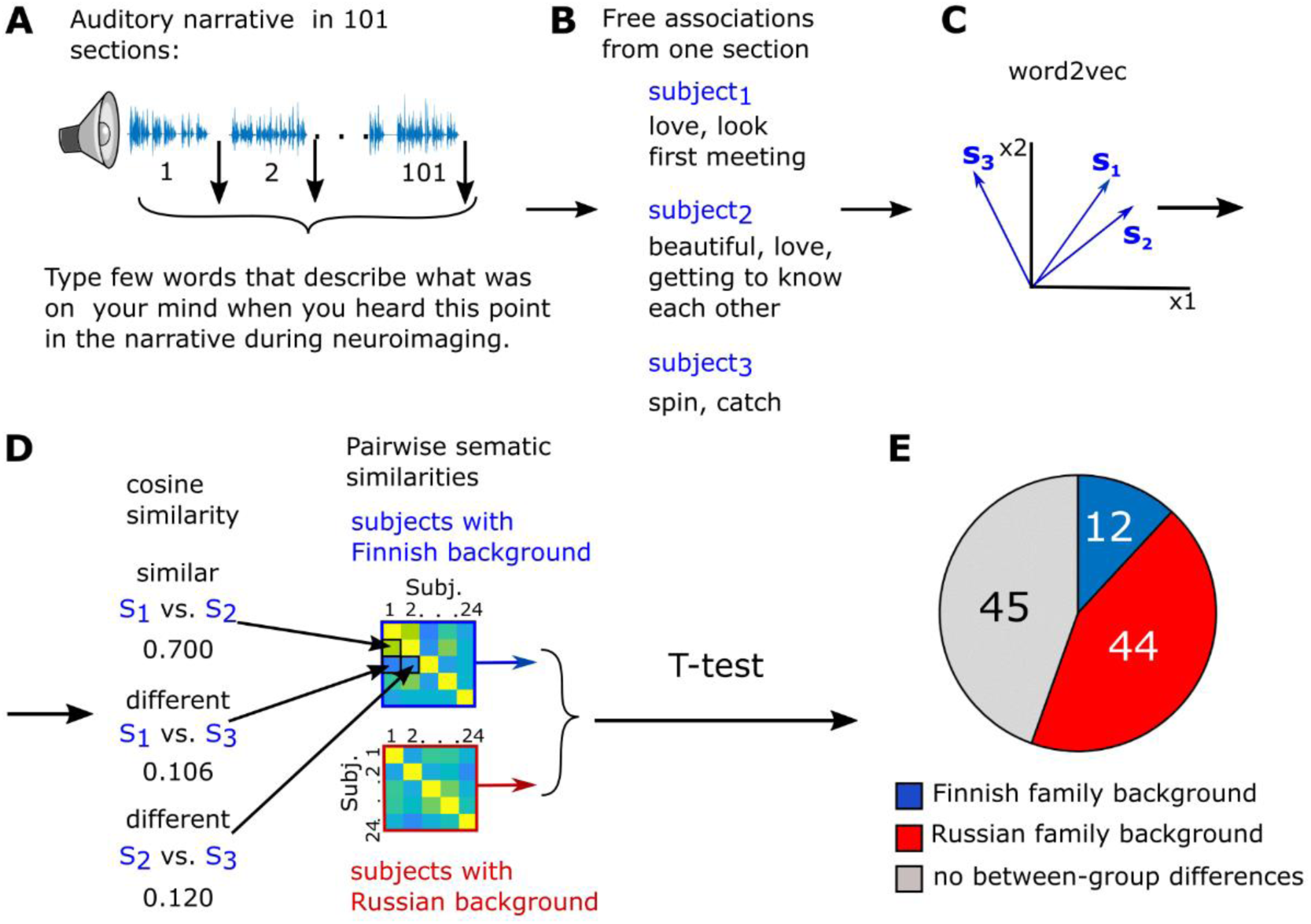
Schematic illustration and results of the behavioral association task. **(A)** In the association task, the audiobook was presented to the subject in 101 segments after the fMRI session. **(B)** After each segment, the subject typed in 20–30 sec words that described what was on his/her mind at that point of the audiobook during fMRI (20). **(C)** The associated words were transformed into the vector representations in a semantic space (word2vec). For each segment, a vector sum was calculated over the vector representations of the words the subject had produced. **(D)** Thereafter, pairwise cosine similarities were calculated between these semantic vectors. Then, t-statistics was used to examine whether the cosine similarities differ between the word lists produced by the two groups of subjects. **(E)** The Finnish-background subjects produced semantically more similar word lists in 12 out of 101 segments, the Russian-background subjects listed semantically more similar words in 44 out of 101 segments. There were no significant effects in 45 segments.

### Cultural family background modulated inter-subject correlation of brain activity during audiobook listening

Audiobook listening elicited significant ISC in several brain regions in both groups (Fig. S1 and Table S1; statistical maps can be found from Neurovault: https://neurovault.org/collections/LINKKSTF/). In the subjects with Finnish *vs*. Russian background stronger ISC was found in the left hemisphere in an area extending from the Heschl’s gyrus (HG) and insula to the superior temporal gyrus (STG) as well as in an area extending from the lingual gyrus (LG) to the middle occipital gyrus (MOG) and cerebellum (Fig. 2, cerebellum not shown; Table S2). In the right hemisphere, ISC was stronger for the Finnish-background subjects in an area including parts of the middle temporal gyrus (MTG), MOG, lateral occipital cortex (LOC) and cerebellum. The ISC was stronger in the Russian *vs*. the Finnish -background subjects in left-hemisphere areas extending from the HG to STG and MTG, as well as bilaterally in posterior-inferior parts of the PCun and anterior parts of the cuneus.

**Fig. 2:**
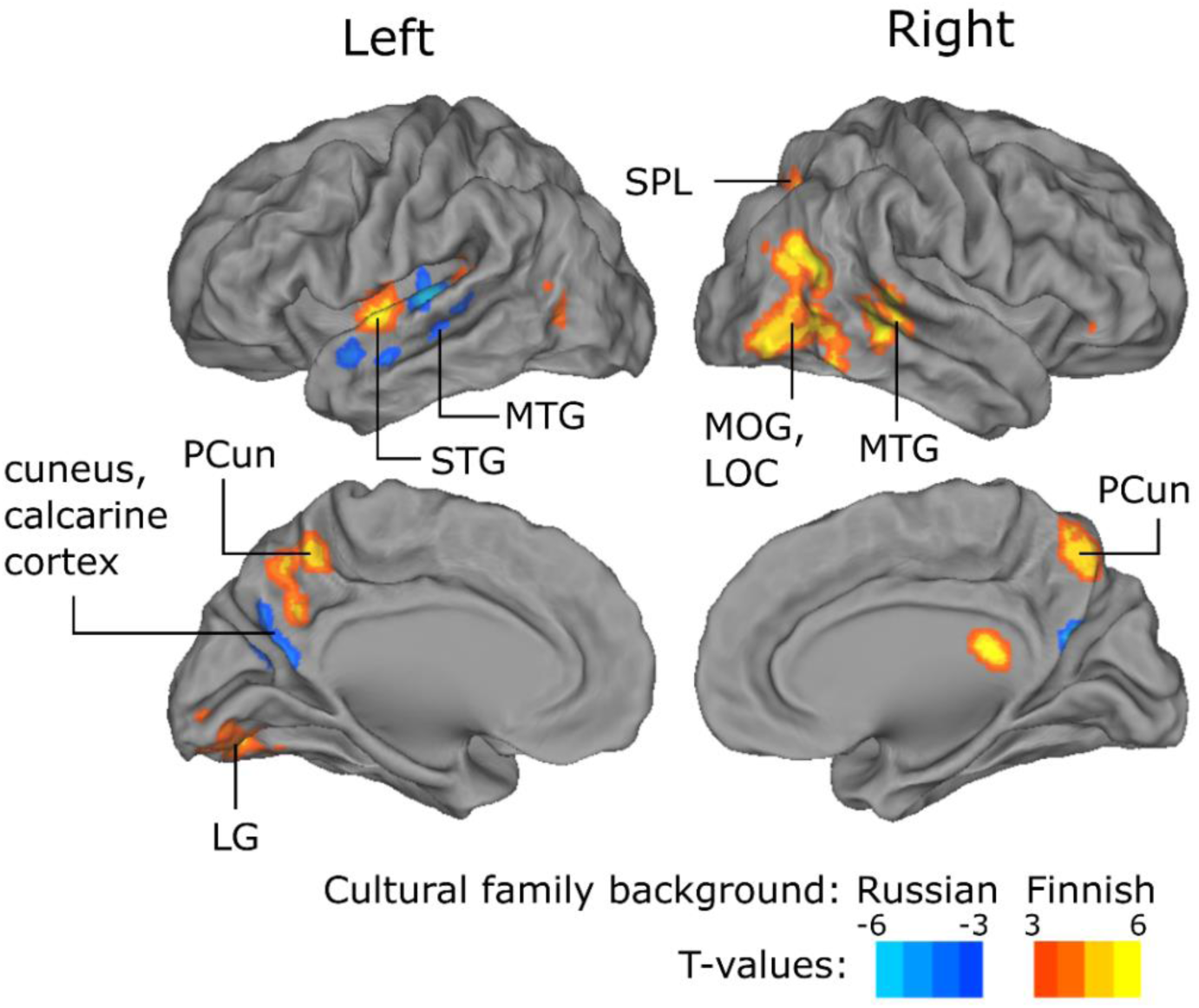
Brain areas where the activity was shaped by family cultural background. The brain areas where ISC was significantly different between the subjects with the Finnish vs. Russian family backgrounds (p<0.05, adjusted with cluster-based thresholding using 5000 permutations and cluster-defining threshold of p < 0.001, uncorrected). Abbreviations: STG = superior temporal gyrus, MTG = middle temporal gyrus, LOC = lateral occipital cortex, MOG = middle occipital gyrus, PCun = precuneus, SPL = superior parietal lobule, LG=lingual gyrus. The loci of ISC were labelled according to the Harvard-Oxford Cortical Structural Atlas (30) implemented in FSL.

### Cultural identification modulated inter-subject correlation of brain activity during audiobook listening

To investigate which of the background variables (Table 1 in Methods and IAT scores in Supplementary materials) are reflected in ISC values, the representational similarity analysis (RSA) was conducted between each of the background similarity matrices and ISC matrix in each voxel. Only the between-subject similarities in the social identification (“how Russian the subject feels herself / himself”) predicted BOLD-similarities between the Russian-background subjects bilaterally in the MTG and PCun as well as in the right angular gyrus (AG, see Fig. 3 and Table S3). We failed to observe significant effects within the Finnish-background subject group, perhaps due to less variability in their social identity self-ratings. The results were obtained using fMRI data that was temporally smoothed with a 2-sec time window (50% overlap). We also studied how the background variables are reflected in the associated words reported in the association experiment. The analysis and results are described in Supplementary text and Fig. S2.

**Table 1.** Results from the background questionnaire. Results are mean values ± SEM. Level of command of Russian language: 1 = I don’t speak Russian, 2 = poor, 3 = satisfactory, 4 = good, 5 = native; Visits in Russia: 1. never visited, 2. visited few times, 3. visited several times, 4. visited regularly, 5. lived longer time periods; How Finnish I feel myself: 0 = not at all, 100 = strongly, How Russian I feel myself: 0 = not at all, 100 = strongly; How positively / negatively I see Finns: 0 = very negatively, 100 = very positively; How positively / negatively I see Russians: 0 = very negatively, 100 = very positively. Significance levels of between-group differences: n.s. > 0.05, *p < 0.05, **p <0.01, ***p<0.001, tested with 50 000 permutations.

| Cultural background | Level of Russian language<br>*** | Visits to Russia<br>*** | How Finnish do I feel myself?<br>(scale 1–100)<br>** | How Russian do I feel myself?<br>(scale 1–100)<br>*** | How positively/negatively do I see Finns<br>(scale 1–100)<br>n.s. | How positively/negatively do I see Russians<br>(scale 1–100)<br>n.s |
| --- | --- | --- | --- | --- | --- | --- |
| Finnish | 1.2±0.1 | 1.4±0.1 | 93.3±1.9 | 1.6±0.3 | 86.3±2.2 | 59.7±2.7 |
| Russian | 4.7±0.1 | 3.9±0.1 | 69.6±3.1 | 55.8±3.2 | 77.3±2.6 | 67.3±2.8 |

**Fig. 3.**
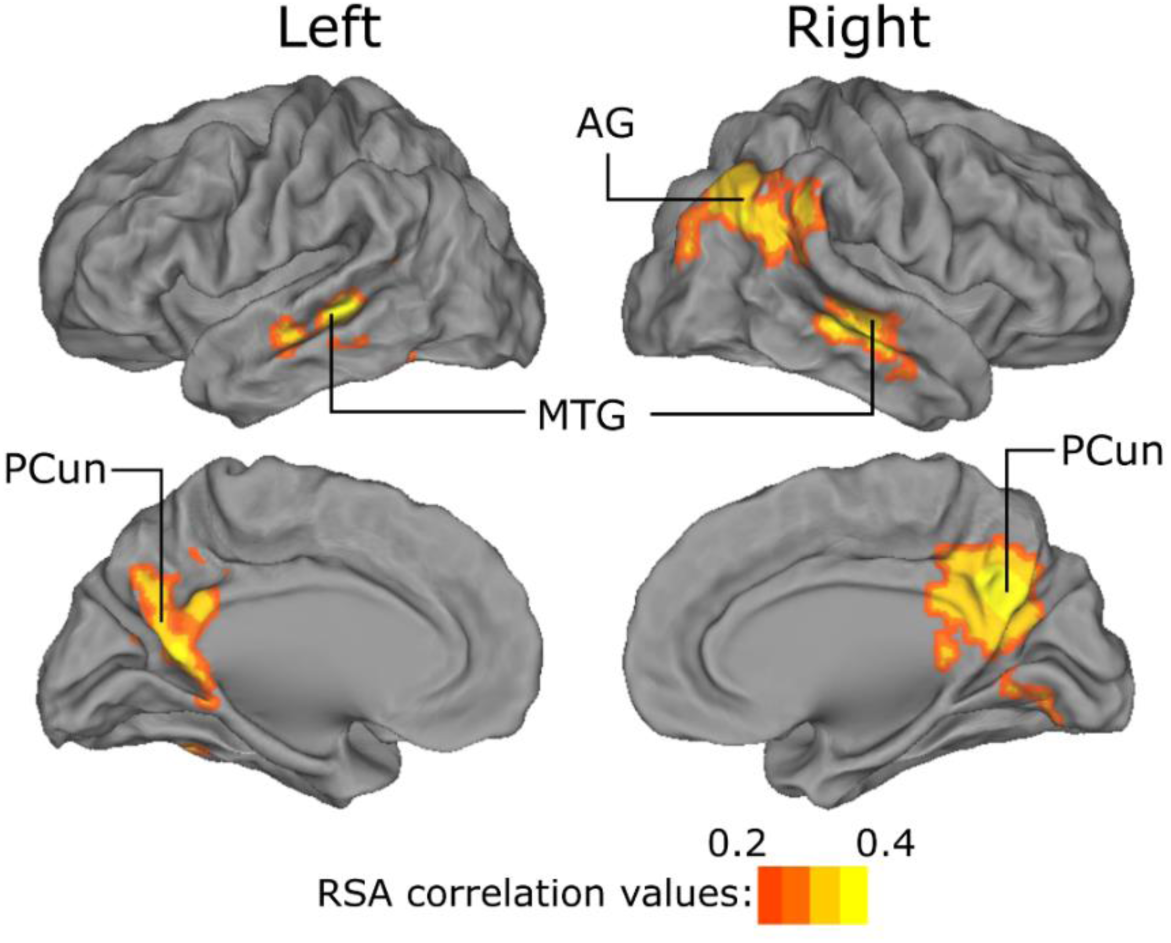
Brain areas where activity was shaped by cultural identification. The brain areas where the between-subject similarities in the question about how Russian the subject feels herself / himself predicted BOLD-similarities between Russian-background subjects (statistical significance tested with 5000 cluster-wise permutations; cluster-defining threshold p < 0.001, uncorrected; the 0.05 family-wise-error corrected cluster size was 261 voxels).

### Implicit association test results

IAT (*29*) was used to investigate whether the subjects unconsciously favored Finland or Russia. Subjects categorized positive and negative adjectives as well as words and pictures related to Finland and Russia while their reaction times were measured (see Supplementary materials for the detailed description). The IAT effect was different between the Finnish- and Russian-background subjects (p<0.001, Fig. S3). Wilcoxon test showed that Finnish-background subjects preferred Finland over Russia (p<0.001, mean d-score: 0.51, std: 0.44, negative values mean preference for Russia and positive for Finland) whereas the Russian-background subjects favored neither country (p = 0.69, mean d-score: −0.07, std: 0.45). Between-group differences in other background variables are shown in Table 1 (Methods).

## Discussion

In the present study, the behavioral word-listing experiment (*20*) disclosed that 56 out of the 101 audiobook segments elicited significantly different associations between the subjects with the Finnish and Russian family backgrounds (Fig. 1), even though all subjects were fluent in Finnish and 44 of had been born and raised in Finland and the remaining four had moved to Finland as small children. The semantic similarity of the associated words was higher among the Russian-background subjects in 44 segments of the story compared to the Finnish-background subjects and vice versa in 12 segments. This suggests that those story segments contained elements that elicit family cultural background related associations specific to each group, and may have been understood more similarly within each group. These segments contained both culture-specific but also non-specific elements (see English-translated text of the audiobook in Supplementary Materials) suggesting that the family cultural background can also modify perception of elements that are not at least obviously related to the culture. Interestingly, out of the cultural background variables 1) the IAT scores, 2) the question how Russian the subjects felt themselves, and 3) the question how Finnish the subjects felt themselves best explained the similarities-differences in the word listings, and for example level of command of Russian language had a negligible effect (Fig. S2, Supplementary text). Overall, the Russian-background subjects produced 33% more words than the Finnish-only background subjects, which could be explained by the Finno-Ugric culture having been observed to be a less talkative one (*31*). Interestingly, then, it seems that the family background can override the general tendency of the mainstream culture towards small talk.

Using fMRI, we were able to reveal, for the first time, that the differences in the family cultural backgrounds of the subjects were reflected as between-group differences in ISC of brain hemodynamic activity in surprisingly many brain regions (Fig. 2), even when both subject groups had equally good command of Finnish and were citizens of Finland. This is an important extension to the previous behavioral studies that have found cross-cultural differences between subjects during story reading as well as neuroimaging studies that have mainly used simple isolated pictures to investigate perception differences of people from Western and Eastern cultures. STG, MTG and PCun have been associated with speech comprehension and processing of semantic information in several previous neuroimaging studies ((*32*), for a review, see (*33*)). Lerner et al. (*15*) showed that words and sentences activated areas in the temporal cortices, and when the sentences formed full paragraphs, the activation extended also to the frontal cortices and PCun. Further, it has been shown that PCun exhibits increased ISC between subjects who have understood visual and auditory narratives more similarly (*25*), is involved in processing paragraphs about in-group *vs*. out-group characters (*34*) and, especially posterior PCun, is also important in episodic memory retrieval (*35*). In the light of these previous studies and the results of the association task, the between-group differences in the ISC in STG, MTG and PCun could be explained by the family cultural background shaping the processing of semantics at multiple timescales as well as in the process of linking the story events into the subject’s prior experiences and memories. Between-group differences in the ISC in the LOC, LG and MOG possibly reflected differences in visual imagery elicited by listening to the audiobook (*36, 37*). The between-group differences in ISC may additionally reflect identifying only with the subjects’ own culture and differential viewing of the protagonists belonging to the subjects’ own *vs*. other cultural group. Indeed, the RSA revealed that out of the cultural background variables, the question how Russian the subjects feel themselves predicted the similarities-differences of the ISC between the Russian-background subjects in the MTG, PCun and AG (Fig. 3., Table S3). Thus, it seems that the sense of belonging to a cultural group, social identity (*38*), shapes the narrative processing in the brain. Interestingly, between-group differences in ISC were also found in the primary auditory cortex at HG. This suggests that the cultural family background modulates even prelexical processing of speech, perhaps via top-down influences from the higher-level speech processing areas.

In conclusion, our results suggest that even a relatively subtle differences in cultural background can result in higher similarity within one group compared to the other in how natural speech is interpreted and how it is processed at multiple hierarchical levels in the human brain. Our results suggest that the strength of social identity most robustly shapes narrative processing in lateral-temporal and temporo-parietal areas that likely process natural speech at word and sentence levels, as well as in precuneus that is known to process natural speech at narrative level. These effects might play an important role in enhancing mutual understanding between individuals with shared cultural backgrounds. Awareness of these effects could help overcome potential challenges in mutual understanding between individuals with different cultural backgrounds and social identities in modern multicultural societies wherein individuals from different backgrounds speak the same native language and might erroneously take for granted highly similar understanding. Enhanced awareness and scientific understanding of these differences in language use is important to understand and avoid obstacles in social interactions at many levels of society.

## Materials and Methods

### Subjects

Forty-eight subjects participated in the study: 24 subjects with Finnish family background and 24 subjects with Russian family background. The subjects with the Finnish family background had Finnish parents and were born and raised in Finland. They were aged between 19–35 years (mean age: 24.7 years). 12 of the Russian-background subjects had both parents Russian, 11 had Russian mother and one had Russian father. The subjects with Russian family background were aged between 18–35 years (mean age: 24.7 years). 20 of the Russian-background subjects were born in Finland, one subject had moved to Finland at the age of eight, one at the age of one, and two at the age of three years. 10 of the subjects with the Russian family background were exposed to both Finnish and Russian in first years of life, 13 started to acquire Finnish at the age of three years when they went to preschool and one at the age of 8 years. All subjects were using Finnish actively in their daily live. Of the subjects with the Russian family background, 19 felt Finnish and two felt Russian as their stronger language, and three felt that both languages are equally strong. The subjects did not constitute a representative sample of Finnish and Russian family background residents of Finland, and therefore, the results cannot be generalized to the whole population with Russian family background people in Finland.

All subjects were right-handed (Edinburgh Handedness Inventory (*39*)). When asked, none of the subjects reported any neurological or audiologic deficits. All subjects were residents of Finland (11 Russian background subjects had also Russian citizenship), fluent in Finnish and had graduated from the Finnish primary school. Half of the subjects in both groups were females. In both groups, one of the subjects had vocational examination and not Matriculation examination and the others had Matriculation examination, undergraduate examination or university degree examination. Prior to participation, all subjects gave their written voluntary informed consent. The study was approved by the Aalto University Research Ethics Committee and conducted in accordance with the Declaration of Helsinki.

### Stimulus

During the fMRI the subjects were presented with a 71-min fictional audiobook in Finnish (custom written by author IPJ; in the end of the Supplementary materials is the English-translated textual version that conveys the events of the story however without artistic level of literacy of a professional translation). The narrative included episodes related to and occasionally contrasting Finnish and Russian cultures (e.g. religion, food, literature, history and traditions). The protagonists of the narrative were two friends, young adults with, Finnish and Russian backgrounds, both living in Helsinki, Finland. In short, the Finnish man was courting a Russian-background woman, whereas the Russian man was in the relationship with a Finnish woman. The characters were described as having personality and behavioral characteristics stereotypically associated with Finns or Russians (e.g. Russian characters were more emotional and conservative than Finns) however without exaggerating to present the protagonists as realistic persons. Cultural differences in expressing emotions, style of thinking, behavioral patterns, values and attitudes resulted in difficulties in understanding each other and in conflicts between the characters. At the end of the story, the Russian-background man ended up in a relationship with the Russian-background woman and the Finnish man with the Finnish woman. The aim of the story was that the Finnish-only background subjects would identify more with the Finnish protagonists and understand the Finnish-culture specific elements, while the subjects with the Russian background would have greater understanding and thus identification with the Russian protagonists than the Finnish-only background subjects and, further, that the culture-specific elements. In the audiobook, periods describing interactions between protagonists were interspersed with periods describing the seasons in the city without social interactions or culture-specific elements. The audiobook was recorded with a sampling frequency of 44 100 Hz in a professional recording studio. For the fMRI experiment, the auditory file of the audiobook was divided into 10 segments and for the association task into 101 segments.

### Experimental procedures

Subjects filled out the background questionnaire (Table 1) as part of behavioral questionnaires. Thereafter, subjects participated in fMRI measurement and simultaneous magnetoencephalography / electroencephalography (MEG / EEG) measurement sessions that were separated by at least one month in order to reduce possible learning effects. The order of the neuroimaging sessions was counterbalanced across the family backgrounds and genders of the subjects. MEG/EEG results will be reported separately.

After the neuroimaging measurements, the subjects performed the IAT which aims at measuring automatic associations or beliefs that the subject is not willing or self-aware to report (*29*). Further, subjects filled an on-line association task at home. In this, the audiobook was re-presented to the subject in 101 segments and at the end of each the subjects were instructed to type a few words that best describe what was on his/her mind at the end of the segment when he/she had heard that segment first time during neuroimaging. The segments are indicated in the English-translated version of the audiobook in the end of the Supplementary Materials.

### Semantic similarity analysis of the self-reported word lists

To examine between-group differences in word lists produced by the subjects in the association task, word2vec skip-gram method (*28*) was first used to transform the individual words into vectors representing semantic-content. Here, the word2vec skip-gram model (Gensim Python Library, https://pypi.org/project/gensim/) was trained using the Finnish Internet Parsebank corpus (*40*) (http://bionlp.utu.fi/finnish-internet-parsebank.html) with 500 dimensions and a window of 10 words. The words that occurred less than 50 times in the corpus were excluded from the model training. The vector embeddings were adjusted by passing the corpus through the word2vec skip-gram model five times.

The model contained 98% of the words produced by the subjects, and the rest of the words were discarded. The word lists produced by the subjects were first corrected for spelling errors and stop-words were removed in the case the subject had written sentences. Thereafter, word2vec was used to map each word to a semantic vector. The semantic representation of each list was obtained by calculating the vector sum over the words in the list. The list vector was computed for 101 narrative segments. Between-subject pairwise semantic similarities were then obtained by calculating cosine distances between the list vectors.

The pairwise cosine similarities within the Finnish-background subjects was compared to the pairwise cosine similarities between Russian-background subjects using a nonparametric T-test where the null-distribution was created by randomly permuting the cosine similarities between groups 50 000 times (*41*). T-test was calculated separately for each of the 101 segments. In addition, the semantic similarities of the associated words across the whole audiobook were estimated by calculating an average over the pairwise cosine similarities over the 101 audiobook segments after weighting the cosine similarities by the length of the segment. Thereafter, the T-test was calculated between the resulting cosine similarities obtained for the subjects with the Finnish and Russian family backgrounds.

### fMRI acquisition

The fMRI session consisted of the following 13 runs: 1) T1-weighted anatomical, 2) a run comprising of four repetitions of the introduction of the narrative (5.93 min), 3) the narrative presented in 10 runs (durations: 4.8–8.4 min), 4) the run comprising of four repetitions of the introduction of the narrative (5.93 min), 5) resting state (3 min) and 6) T2-weighted anatomical. Each functional run started with a period of 12.3 s and ended with a period of 15 s without auditory stimulation. The repetitions of the introduction were separated by a period of 16 s without auditory stimulation. The subject was shown a white fixation cross on a black background during the functional runs (including resting state) and images of Helsinki between the runs and during the measurement of reference scans. 24 subjects wanted to have one, six subjects 2–4, and 18 subjects zero breaks outside of the scanner between runs.

Anatomical and functional MRI data were acquired with a 3T MRI whole-body scanner (MAGNETOM Skyra, Siemens Healthcare, Erlangen, Germany) using a 32-channel receiving head coil array. In the beginning of the measurement session, anatomical images were measured using a *T*_1_-weighted MPRAGE sequence (TR=2530 ms, TE = 3.30 ms, field of view (FOV) = 256 mm, flip angle = 7 degrees, slice thickness = 1 mm). Whole-brain fMRI data was measured using an ultra-fast simultaneous multislice (SMS) inverse imaging (InI) sequence (*27*). The SMSInI has higher spatial resolution with lower signal leakage and higher time-domain signal-to-noise ratio than inverse imaging without SMS, and detects subcortical fMRI signals with similar sensitivity and localization accuracy as echo planar imaging (*27*). Here, the SMSInI enabled more accurate removal of physiological artifacts (*42*) and utilization of the higher sampling rate under conditions of listening to natural connected speech wherein words occur at rates of ∼three per second. Instead of completely relying only gradient coils in spatial encoding, InI achieves spatial encoding by solving the inverse problems utilizing the spatial information from channels in a radio-frequency coil array and gradient coils. In this study, InI-encoding direction was superior-inferior, whereas frequency and phase encoding were used to recover the spatial information in anterior-posterior and left-right directions, respectively. 24 axial slices (7 mm) were first collected without gap between slices. Thereafter, the slices were divided into two groups of 12 slices, and each of the slice groups was excited and read in 50 ms resulting in a TR of 100 ms. Simultaneous echo refocusing (*43*) was used to separate adjacent slices in each group, and aliasing was further controlled with blipped controlled aliasing in parallel imaging (*44*). Other measurement parameters were: TE=27.5 ms, flip angle=30°, FOV=210 × 210 × 210 *mm*^3^, and in-plane resolution = 5 mm × 5 mm.

Solving an inverse problem in InI reconstruction requires sensitivity map of the channels in the coil array (*27*). This information was included in a 6-sec reference scan measured before each functional run. In the reference scan, partition-encoding steps were added after slice group excitation in InI-encoding direction. The reference scan and accelerated scans were acquired with the same other imaging parameters. Before each reference scan, shimming was used to minimize inhomogeneity in the magnetic field.

During the fMRI measurements, stimulus presentation was controlled with Presentation software (Neurobehavioral Systems, Albany, NY, USA). The auditory stimuli were presented to the subject through Klaus A. Riederer (KAR) ADU2a insert earphones. The intensity level of the stimulus was adjusted on an individual subject basis at a comfortable listening level that was clearly audible above the scanner noise. The audio out from the sound card of a computer was recorded with the BIOPAC MP150 Acquisition System (BIOPAC System, Inc.). In the data analysis, this allowed us to determine the exact times when the auditory stimulus has started. The BIOPAC MP150 system was also used to record heart rate and respiration signals during the fMRI measurement. Heart rate was measured using two BIOPAC TSD200 pulse plethysmogram transducers placed on the palmar surfaces of the subject’s left and right index fingers. Respiratory movements were measured using a respiratory-effort BIOPAC TSD201 transducer attached to an elastic respiratory belt, which was placed around the subject’s chest. Heart rate and respiratory signals were sampled simultaneously at 1 kHz using RSO100C and PPG100C amplifiers, respectively, and BIOPAC AcqKnowledge software (version 4.1.1).

### fMRI reconstruction and preprocessing

Anatomical images were reconstructed using Freesurfer’s automatic reconstruction tool (recon - all; http://surfer.nmr.mgh.harvard.edu/) and functional images using the regularized sensitivity encoding (SENSE) algorithm with a regularization parameter of 0.005 (*45, 46*). The reconstructed images were registered to the Montreal Neurological Institute 152 (MNI152) standard space template by first calculating transformation parameters from structural to standard space and from the reference scan to structural space. Thereafter, these transformations were concatenated and used to co-register functional images to the MNI152 standard space with 3-mm resolution. The co-registrations were performed by Freesurfer and FSL tools (*47, 48*). A period of 12.3-s of fMRI data measured before the start of the audiobook was removed from each fMRI measurement. To remove the scanner drift, the data was detrended using a Savizky-Golay filter (order: 3, frame length: 240s). Physiological and movement artifacts were suppressed using MaxCorr method (*49*). Specifically, from the data of each subject, we regressed out ten components that correlated maximally within the white matter and cerebrospinal fluid of that subject but were minimal in the other subjects’ white matter and cerebrospinal fluid. Since these components were subject-specific, they were assumed be to the artifacts rather than brain activity elicited by the audiobook. Thereafter, DVARS (*50*) was used to identify the data potentially affected by head motions. No differences were found in DVARS values between the two subject groups suggesting that the between-group differences found in the ISC values should not be related to the head movements (Finnish family background: 1.64±0.46, Russian family background: 1.59 ±0.41, T-value: 0.70, p=ns., nonparametric T-test with 5000 permutations). The fMRI data was filtered between 0.08 and 4 Hz using a zero-phase filter and smoothed spatially with a 6-mm full-width-half-maximum Gaussian kernel.

### Inter-subject correlation (ISC) analysis of BOLD-responses

BOLD-responses between subjects were compared using ISC analysis as implemented in the ISC-toolbox (https://www.nitrc.org/projects/isc-toolbox/) (*51*). For each voxel and for each of the 10 runs, Pearson’s correlation coefficients were calculated across the time-courses of every subject pair, resulting in 1128 (*n* × (*n*− 1)/2, where n = 48) ISC values per voxel. To calculate ISC values across the whole audiobook, average ISC values were calculated over the run-wise ISC values that were first transformed to z-values using Fisher’s transformation and weighted with the length of the run. The statistical significance of the ISC values was evaluated by a nonparametric voxel-wise resampling test to account for the temporal autocorrelations in the BOLD data (*51*). In short, a null-distribution was created from ISC values calculated after circularly shifting the time series for each subject by a random amount such that the timeseries between subjects became unaligned in time. The null-distribution was approximated with 0.5 million realizations randomized across voxels and time-points. The null-distribution across the whole audiobook was determined by taking a weighted average over the ten run-wise null-distributions. The threshold for significant ISC was determined by first computing p-values for the true realizations for each voxel based on the null-distribution and, thereafter, correcting the resulting p-values for multiple comparisons using the FDR-correction. Between-group differences of Fischer’s z-transformed ISC values (z-scores) were studied using a nonparametric cluster-based two-sample T-test where the statistical significance was tested by 5000 random permutations of the z-scores between the two groups (*41, 52, 53*). The null-distribution was created from the maximum cluster sizes obtained by thresholding the statistical images at the cluster-defining threshold of 0.001 at each permutation. Corrected p-value for each suprathreshold cluster was obtained by comparing its size to the permutation distribution.

### Background variables which most accurately predict the ISC between subjects

The voxel-wise between-subjects RSA (*54*) was used to test in which brain regions greater similarity in background variables between pairs of subjects predicts greater BOLD-similarity between pairs of subjects. First, ISC matrices were calculated separately between the Finish and Russian family background subjects using temporally smoothed fMRI data (time window of 2 sec, 50% overlap). Next, the corresponding similarity matrices were created for each background question using the score differences between each subject pair. Third, the correlation was calculated between each of the background similarity matrices and BOLD-similarity matrix in each voxel. The statistical significance of the results was tested using a cluster-wise non-parametric permutation test (*52*) with 5000 subject-wise permutations (cluster-defining threshold p < 0.001, uncorrected; the 0.05 family-wise-error corrected cluster size was 261 voxels).

## Acknowledgments

This research was financially supported by the Academy of Finland (grant No. 257811, 273469, 276643 and 287474), Jane and Aatos Erkko Foundation, Emil Aaltonen Foundation, and Russian Federation Government (grant ag. No. 075-15-2019-1930). The calculations presented above were performed using computer resources within the Aalto University School of Science “Science-IT” project. We also want to thank Dr. Toni Auranen for an excellent technical support in setting up the SMSInI sequence in the AMI centre, Julia Bethwaite for her valuable feedback on Russian cultural aspects of the story, Esko Salervo for voice-fitting and speaking the story, Jenna Kanerva and Filip Ginter at the University of Turku for development of the Finnish language Word2vec model and all the subjects for making this study possible.

## Author contributions

M.H.: Conceptualization, Methodology, Software, Validation, Formal analysis, Investigation, Resources, Data Curation, Writing - Original Draft, Visualization, Project administration, Funding acquisition. A.I.: Methodology, Software, Validation, Formal analysis, Data Curation, Writing - Review & Editing. A.H.: Methodology, Writing - Review & Editing. J.K.: Methodology, Software, Writing - Review & Editing. M.K.: Conceptualization, Methodology, Writing - Review & Editing, Supervision, Project administration, Funding acquisition. F-H.L.: Methodology, Software, Writing - Review & Editing, Funding acquisition. A.L.: Methodology, Resources, Writing - Review & Editing. M.S.: Conceptualization, Methodology, Writing - Review & Editing. I.P.J: Conceptualization, Methodology, Resources, Writing - Review & Editing, Supervision, Project administration, Funding acquisition.

## Competing interests

Authors declare no competing interests.

## Data availability

Stimulus material and codes used in the current study are available from the corresponding author on reasonable request. Pseudonymized fMRI are available within European Union (EU) from the corresponding author by reasonable request in respect of the privacy of the subjects following the guidelines of the Data Protection Act of Finland (includes EU’s General Data Protection Regulation, GDPR). Since the facial features of the anatomical T1 MRI images are needed in the analysis of magnetoencephalography data also measured in this study, T1 images will be available within EU by reasonable request following the guidelines of the Data Protection Act of Finland only after the end of the project when facial features of the MRI volumes can be removed.

## Notes

https://neurovault.org/collections/LINKKSTF/

### Competing Interest Statement

The authors have declared no competing interest.

