## supplementary_information for "Processing of a spoken narrative in the human brain is shaped by family cultural background"

Maria Hakonen, Arsi Ikäheimonen, Annika Hultèn, Janne Kauttonen, Miika Koskinen, Fa-Hsuan Lin, Anastasia Lowe, Mikko Sams, Iiro P. Jääskelainen

**This PDF file includes:**

Supplementary Text
Figs. S1 to S3
Tables S1 to S3
References
English-translated text of the audiobook

### Supplementary text

#### Background variables which most accurately predict the association experiment results

We used boosted decision tree regression to test which of the background variables (Table 1 and IAT scores, Fig. S3) would most accurately predict the interpretations of the narrative in the association experiment. The boosted decision tree regression is a nonlinear regression method that also allows determining the importance of variables. First, we created a between-subject similarity matrix for each background question using the score differences between each subject pair. This resulted in data matrix of 1128 values ($n \times (n-1)/2$, where $n$ = 48; 24 rows and 24 columns). Then we used 10 randomized cross-validation folds and fitted the regression model for each fold using 90% of the data for training and 10% for testing. We used MATLAB's function "fitrensemble" (default parameters) and Bayesian hyperparameter optimization to find optimal tree regressors for the training data. Predicted responses for the testing data were then compared to the actual values. Feature importance weights were obtained from each of the ten models. The weights correspond to the relative importance of the background variables in predicting the semantic similarities of the associated words. The mean importance values and one standard deviation are depicted in Fig. S2. The correlation between real and predicted values over 10 folds was 0.052.

#### Implicit Association test (IAT)

The IAT comprised of seven tasks. The data was analyzed from the fourth and seventh tasks, and the other tasks introduced the IAT to the subject. In the first task, images and words related to Finnish or Russian cultures appeared in the middle of the computer screen and the two categories "Finland" and "Russia" were shown in the left and right corner of the screen, respectively. The subject was instructed to categorize the presented items into the appropriate category as quickly as possible by pressing the appropriate left-hand or right-hand key. The second task was to categorize positive and negative adjectives into the categories “Good” (left corner) and “Bad” (right corner) with the same procedure as in the first task. In the third task, the previous categorizes were combined into two categories “Finland and good” (left corner) and “Russia and bad” (right corner), and the subject was instructed to categorize the examples from all four categories into the appropriate combined category. The fourth task was a repeat of the third task. The fifth task was similar as the first task with an exception that the position of the names of the categories was reversed (i.e. left corner: “Russia”, right corner: “Finland”). The sixth task was similar as the third one, except that the pairings of the categories was opposite (i.e. left corner: “Russia and good”, right corner: “Finland and bad”). The final task was a repeat of the sixth task. The task order of the IAT was counterbalanced across the family background and gender of the subjects such that for one half of the subjects the positions of the categories “Finland” and “Russia” were reserved in all tasks (i.e. Russia was first paired with good and Finland with bad). The subject was obligated to correct errors before proceeding, and reaction times were measured to the occurrence of the correct response. The IAT effect was calculated for each subject applying the Choen’s d to the response latencies from the fourth and seventh tasks as follows

$$d = \frac{Fi/BAD - Fi/GOOD}{std_{Fi/BAD-Fi/GOOD}} + \frac{Ru/GOOD - Ru/BAD}{std_{Ru/GOOD-Ru/BAD}}$$

where $Fi/BAD$ is the mean latency for the combined category “Finland and bad”, $Fi/GOOD$ the mean latency for the combined category “Finland and good”, $Ru/GOOD$ the mean latency for the combined category “Russia and good”, $Ru/BAD$ the mean latency for the combined category “Russia and bad”, $std_{Fi/BAD-Fi/GOOD}$ the standard deviation of the latencies for the categorizes “Finland and bad” and “Finland and good” and $std_{Ru/GOOD-Ru/BAD}$ the standard deviation of the latencies for the categorizes “Russia and bad” and “Russia and good”. The more positive the IAT effect was, the more the subject was interpreted to prefer Finland over Russia. The more negative the IAT effect was, the more the subject was interpreted to prefer Russia over Finland. A nonparametric two-sample T-test with 5000 permutations was used to examine whether the IAT effect was different between the two groups. Wilcoxon signed rank test was applied separately to the d-scores of the Finnish-background and Russia-background subjects to examine whether the subjects within either of the groups preferred Finland over Russia or vice versa.

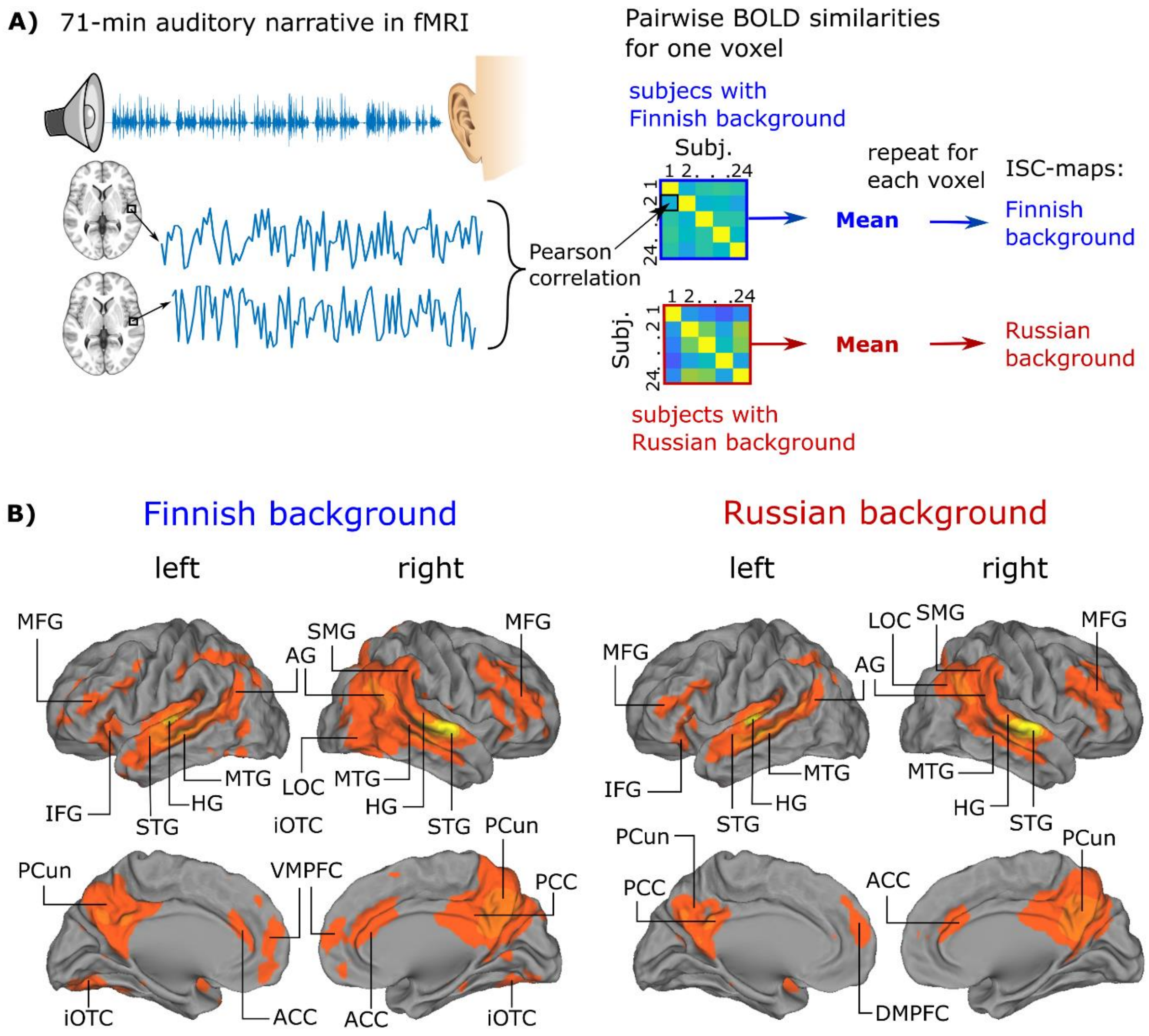


***Fig. S1. ISC during narrative listening. (A)*** *Schematic illustration of the experimental paradigm and ISC analysis (see Methods for details).* ***(B)*** *Significant ISC of hemodynamic activity displayed on cortical surfaces at a voxelwise false-discovery rate (FDR) threshold of 0.001. Abbreviations: MFG = middle frontal gyrus, AG = angular gyrus, IFG = inferior frontal gyrus, STG = superior temporal gyrus, HG = Heschl's gyrus, MTG = middle temporal gyrus, PCun = precuneus, iOTC = inferior occipitotemporal cortex, LOC = lateral occipital cortex, VMPFC = ventromedial prefrontal cortex, ACC = anterior cinculate cortex, PCC = posterior temporal cortex, DMPFC = dorsomedial prefrontal cortex. The loci of ISC were labelled according to the Harvard-Oxford Cortical Structural Atlas implemented in FSL.*

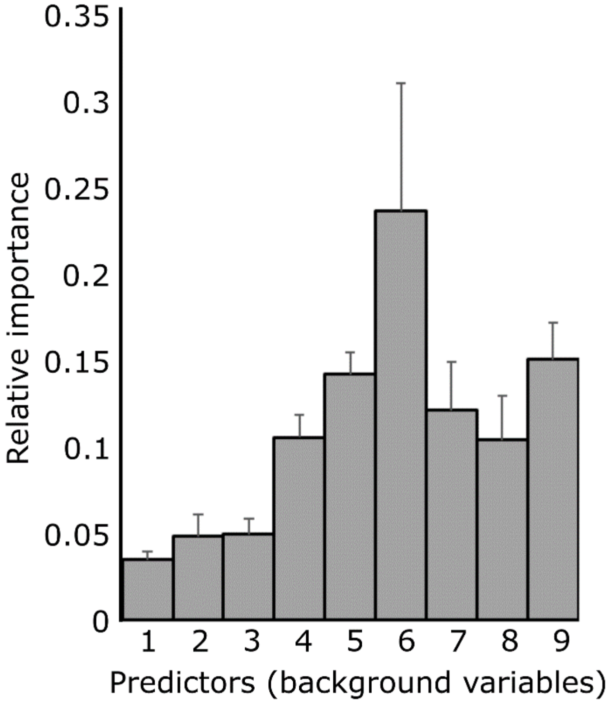


***Fig. S2. Relative importance of background variables in predicting between-subject similarities in the association experiment****. Relative importance values are means over 10 folds and error bars describe the standard error of the relative importance values over 10 folds. 1 = nationality (Finnish or Russian), 2 = Number of Russian parents, 3 = The level of Russian language, 4 = How often do I visit Russia, 5 = How Finnish do I feel myself, 6 = How Russian do I feel myself, 7 = How positively / negatively do I see Finns, 8 = How positively / negatively do I see Russians, 9 = IAT score.*

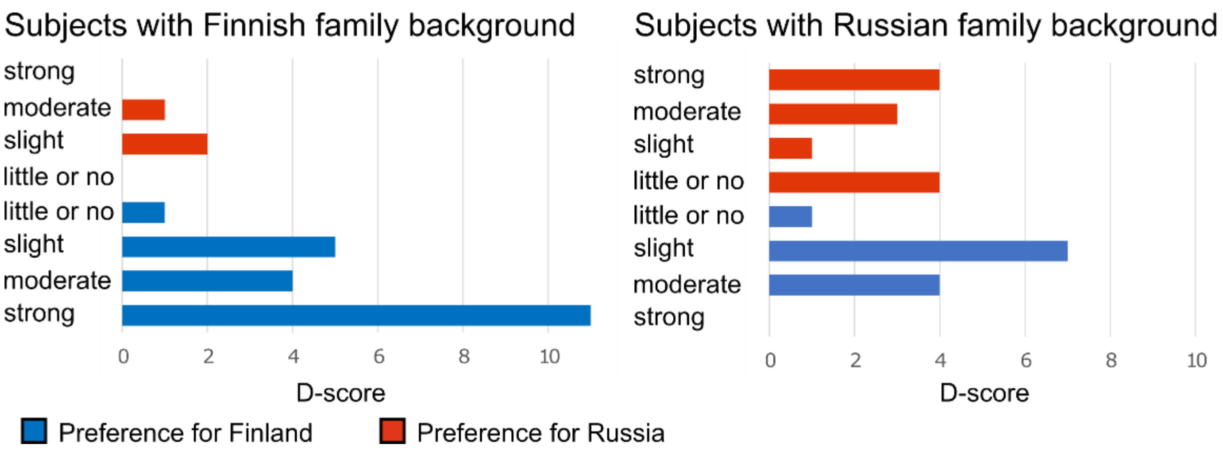


***Fig. S3. Subjects' preferences for Finland and Russia obtained from IAT.*** *Bar graphs present the number of the subjects with each preference level. D-score threshold values were as follows: ≤ -0.65: strong preference for Russia over Finland, >-0.65 and ≤ -0.35: moderate preference for Russia over Finland, > -0.35 and ≤ -0.15: slight preference for Russia over Finland, > -0.15 and < 0.15: little to no preference between Finland and Russia, ≥ 0.15 and < 0.35: slight preference for Finland over Russia ≥ 0.35 and < 0.65: moderate preference for Finland over Russia, ≥ 0.65 strong preference for Finland over Russia* (*1*, *2*).

***Table S1. ISC values across Finnish- and Russian-family-background subjects.*** *Labels, sizes, peak coordinates, and maximum ISC-values of clusters obtained in the ISC-analysis for the subjects with Finnish and for the subjects with the Russian family backgrounds. The loci of ISC were labelled according to the Harvard-Oxford Cortical Structural Atlas implemented in FSL.*

| Cluster label | Cluster extent (voxels) | x MNI (mm) | y MNI (mm) | z MNI (mm) | max ISC value |
|---|---|---|---|---|---|
| **ISC Finnish background** | | | | | |
| Planum temporale (R) | 1730 | 60 | -12 | 0 | 0.14 |
| Heschl's gyrus (L) | 1568 | -48 | -18 | 3 | 0.14 |
| Frontal Orbital cortex (L) | 82 | -39 | 27 | -9 | 0.02 |
| Frontal pole (R) | 69 | 48 | 42 | 3 | 0.02 |
| Precuneus cortex (R) | 1338 | 6 | -63 | 33 | 0.06 |
| Frontal pole (L) | 68 | -36 | 36 | 9 | 0.02 |
| Cingulate gyrus (R) | 71 | 6 | 33 | 21 | 0.02 |
| **ISC Russian background** | | | | | |
| Cerebellar crus II (R) | 38 | 30 | -75 | -39 | 0.04 |
| Superior temporal gyrus, posterior division (R) | 1055 | 63 | -12 | 0 | 0.14 |
| Heschl's gyrus (L) | 880 | -48 | -21 | 3 | 0.16 |
| Inferior frontal gyrus (R) | 72 | 45 | 18 | 24 | 0.03 |
| Precuneus cortex | 920 | 6 | -63 | 30 | 0.06 |

***Table S2. Differences of ISC values between Finnish- and Russian-family-background subjects.*** *Cluster size, peak coordinates, the maximum T-values and p-values of the clusters obtained in the nonparametric two sample T-test between the two subject groups (5000 cluster-wise permutations; cluster-defining threshold, uncorrected: p < 0.001; the 0.05 family-wise-error corrected cluster size: 62 voxels). The loci of ISC were labelled according to the Harvard-Oxford Cortical Structural Atlas* (*3*) *implemented in FSL.*

| Cluster label | Cluster extent (voxels) | x MNI (mm) | y MNI (mm) | z MNI (mm) | max T-value | p-value |
|---|---|---|---|---|---|---|
| **ISC Finnish > ISC Russian** | | | | | | |
| Temporal pole (superior, R) | 87 | 54 | 21 | -24 | 6.79 | p < 0.03 |
| Middle occipital gyrus (L) | 468 | -42 | -66 | 3 | 6.81 | p < 0.002 |
| Middle temporal gyrus (R) | 481 | 57 | -63 | 12 | 6.92 | p < 0.002 |
| Middle temporal gyrus (R) | 143 | 51 | -42 | 3 | 5.89 | p < 0.02 |
| Superior temporal gyrus (L) | 217 | -57 | -6 | 0 | 6.32 | p < 0.01 |
| Precuneus cortex (R) | 401 | 18 | -60 | 33 | 7.74 | p < 0.002 |
| **ISC Russian > ISC Finnish** | | | | | | |
| Superior temporal gyrus (L) | 210 | -48 | -24 | 6 | -6.25 | p < 0.01 |
| Calcarine cortex (L) | 85 | -9 | -69 | 18 | -5.23 | p < 0.03 |

***Table S3. Correlations between ISC-values and between-subject similarities in identification to Russian culture within Russian-background subjects****. Cluster size, peak coordinates, the maximum correlation values and p-values (5000 cluster-wise permutations; cluster-defining threshold, uncorrected: p < 0.001; the 0.05 family-wise-error corrected cluster size: 261 voxels) of the clusters obtained in the RSA between BOLD-similarities and similarities in the question about how Russian the subject felt herself / himself. The results were obtained for the Russian-background subjects. The loci of ISC were labelled according to the Harvard-Oxford Cortical Structural Atlas (*3*) implemented in FSL.*

| Cluster label | Cluster extent (voxels) | x MNI (mm) | y MNI (mm) | z MNI (mm) | max correlation value | p-value |
|---|---|---|---|---|---|---|
| Middle temporal gyrus (L) | 340 | -45 | -39 | -3 | 0.42 | < 0.04 |
| Middle temporal gyrus (R) | 283 | 51 | -27 | -6 | 0.40 | < 0.05 |
| Precuneus (R) | 984 | 12 | -54 | 33 | 0.35 | < 0.004 |
| Angular gyrus (R) | 426 | 45 | -63 | 30 | 0.31 | < 0.03 |

**English-translated text of the audiobook**

**Blue = similarity higher amongst Finnish-background subjects**

**Red = similarity higher amongst Russian-background subjects**

**Black = no differences in similarity**

Part 1

The sky is like a thick gray carpet, through which the sun glows weakly. Its light has already begun to dim - the evening is coming. Eventually the dark descends and covers the city.

A breezy wind from the sea typically blows in the streets and parks of the city, but now the damp air feels repulsive and sticky. It hasn't rained all day. It is as if someone wanted to delay the eruption of clouds.

Brightly colored, Jugend-era houses and yellow-red leaves that have fallen from the trees bring color to the gloomy autumn view, but when darkness of the eve fall, their colors also blend in with shades of gray.

It is probably due to this depressing weather that the streets of the city center are unusually quiet, even though it's Friday night. It's like everyone's gone.

Part 2

However, there is life behind the countless windows of the densely populated city center. The warm and cozy apartments have a delightful Friday night celebration in full swing. This is also the case in the stylishly modernized one-bedroom apartment located on the third floor of a Jugend house.

"Via Ostrobothnia," Ville, a twenty-year-old slender-build and athletic blonde guy, says, lifting a snaps glass filled with cold Koskenkorva vodka at a small kitchen table by his friend Sasha. The dark-haired and blue-eyed Sasha happily replies, singing, "via Bothnia, via Bothnia". His eyes twinkled merrily. The buddies gulp down the throat-soaring drinks and laugh afterwards.

Part 3

"Let's see if we find good women tonight," Ville says, his thoughts are already moving forward to Helsinki's nightlife.

"Let's see - maybe you meet a woman tonight," Sasha laughs. He bends a little forward and his face spreads into a lively smile. “I have now met a couple of times a really great blonde, Laura. Maybe I don't need to meet any other anymore. ”

Part 4

This knowledge surprises Ville. He doesn't really know what to think. To cover up his confusion, he starts to fool around, first playing astonished and then joking with his friend: "It can't be true, the womanizer Sasha has found someone who is sufficient for him." He fills both vodka shot glasses. "For your new happiness and my eternal search," Ville utters with a bit of a drunken voice. He empties the snap glass with a single shot.

Part 5

For a moment, Ville's gaze is fixed on a decorative icon hanging on Sasha’s kitchen wall. It is as if the mother-Maria with her halo looks at the boys with a motherly disapproving look with the baby in her arms. Ville points with his left forefinger at the icon and giggles: "Look, Maria watches over our drinking behavior", but adds almost instantly: "Yeah bad joke, I did not mean that really" when he notices that Sasha’s smile is a forced one. Ville's apology raises a genuine warm smile on Sasha's face. He cheerfully gulps down the vodka in his glass, grinning.

Part 6

Sasha and Ville move to the sauna and continue to their joyful getting drunk amidst the steam. Now they drink beer. Sasha cannot help telling about Laura one praise after another. Ville listens silently. He is on one hand joyful for his friend's happiness, on the other hand a little jealous. "You've got a lot of luck with ladies,"

Ville finally says, scoops water from a bucket and throws it on the sizzling hot stove. "Well, for a very long time I did not have any luck," Sasha defends and continues, "Like with that one Emilia." Ville bursts into uncontrollable laughter to which Sasha joins.

Part 7

"Sorry when I laughed at it, but it was such a sorry occurrence," Ville says after he stops laughing. "Well – but I had my thing with that Svetlana," Ville continues, and this time Sasha is the one who laughs first. “You were out of sync with Svetlana in just two hours! You don't understand more about a Russian woman than a pig understands with a silver spoon”, Sasha laughs out loud with drunken voice. Ville starts to also laugh. He feels that Sasha's analysis fits perfectly.

Part 8

Sitting in the sauna, Sasha cannot help telling how he got to know Laura. Earlier that week, Sasha had been in a home party. There his gaze met that of a blonde woman. Sasha felt as if electrical current had passed through his body and noticed how the expression of the blond-haired beauty became serious. In the course of  the evening, their gazes met several times while Sasha and the beauty were both engaged talking with others in the party.

Part 9

Eventually Sasha got the opportunity to introduce himself. The blond-haired woman happened to be in the kitchen as Sasha stepped in looking for some more snacks. Sasha felt a lump in his throat. His heart began to beat strongly in the midst of awkward tension induced by shyness in the presence of such a breathtaking beauty. However, Sasha regained his self-confidence when he noticed that the woman blushed when she noticed him. "I am not a loser," Sasha reminded himself and forced himself to breathe in and out slowly. At the same time, he conjured a warm smile on his face. "Here in the kitchen it is less packed than out there in the living room”, Sasha gathered his courage to say. His voice initially trembled slightly, but he managed to steady his voice already before the sentence was over. He continued to confidently say, "How did you find yourself in this party? I'm Sasha. What is your name?".

Part 10

The name of the beauty was Laura. She was a friend of the hostess of the party. Sasha asked Laura to tell her about her studies. Laura began to talk enthusiastically about consumer protection laws. Sasha was generally not interested in this type of thing, but since it was Laura telling, even otherwise boring legislative issues seemed fascinating.

"The rest you will probably guess," Sasha says to Ville, laughing happily and dripping water on the stove.

Ville guesses, of course. For a moment, she feels strange lonely and vulnerable.

Part 11

After the fall of the autumn eve, a dense fog has risen from the sea. There is scent of autumn in the air. There are lonely characters here on the streets of the city. Most have a four-legged friend with them. The characters stop at regular intervals, when interesting odors attract the attention of the hairy companion.

There are, amongst the apartment buildings with metal-sheeted-roofs, green parks, dog-parks, children's playgrounds, old oak trees and lawn areas. Between the old stone houses, lights of passing cars occasionally cut through the darkness, engines whir smoothly, and tires vibrate on cobblestone.

From the sea, a damp, given the season unusually warm, wind begins to rise. It wipes out the stagnant atmosphere and brings with it the smell of salt. The atmosphere is now completely changing. Although the summer's bright nights are now just a memory, the darkness of autumn has not yet succeeded in depressing people's minds. The steps of the people on the streets turn lighter with the wind. People covertly and curiously glance others who they meet on the streets.

From amidst tree branches, jays are cawing. At times, they tend to flock densely over the rooftops, only to return a moment later to the leafless branches of the trees in the park. Colorful leaves cover most of the ground, even though fallen leaves have been raked away on a couple of occasions in the course of the autumn.

Part 12

Fresh from the sauna, Ville and Sasha swing around the corner, with their steps already staggering. Sasha cannot help in his little bit drunken state talking  about his new lovely girlfriend. Ville listens with interest and is happy for Sasha, yet at the same time a bit jealous. He hopes that one day,  sooner rather than later, he will also be blessed with a wonderful girlfriend.

Part 13

The friends are aiming at a popular nightclub at the edge of a park. From afar, they can see that there a 20 meter queue outside the nightclub door. “We’re lucky it is not cold, it would be miserable to queue,” Ville wonders, “you think it’ll take longer than half an hour to get in?” After they’ve taken a few more staggering steps towards the nightclub, Sasha exclaims “It is not necessary to queue, Laura is there right at the door already! “ He lowers his voice and says “lets cut in line to her so we do not need to queue”.

Part 14

The buddies straighten their steps so that the doorman will not find them drunk, and head straight for the beginning of the line. Behind them there are a few drunken protests. Sasha catches Laura from behind to his embrace. Laura is initially startled, but then happily surprised to see that she will get to spend a nice evening with her new boyfriend.

Part 15

Laura is wearing a light long coat with a spectacular jacket underneath. Laura also has high-heeled blond boots that capture the gazes of men.

Laura is accompanied by a dark-haired female friend in blue jeans and a leather jacket. He doesn't seem to feel very happy about the appearance of the boys. She would have liked to chat and have fun amongst ladies. However, he does not want to spoil the good Friday night mood and conjures a friendly smile on his face.

Part 16

"This is my lovely girlfriend Laura," Sasha proudly presents to Ville. "Oh, it is so nice to meet you! Sasha speaks of you all the time," Laura says with a soft voice, shaking hands with him. Ville realizes that he is blushing, but he manages to say with a steady voice that Sasha has also been talking a lot about Laura.

Part 17

That is not quite true – Sasha has just praised Laura's appearance and told Ville how they had met.

"Nice to hear," Laura replies. She looks at Sasha in love and falls again into his embrace.

Part 18

Villen's attention is drawn to Laura's friend, who has long dark curly hair, big brown eyes with slim and relatively tall body. "Oh, you must be Laura's friend. I'm Ville, Sasha's friend," Ville presents with his hand out. "Hi, I'm Olga. It's fun to get to know," the dark beauty responds with Russian accent. "Are you Russian?" Ville asks in a literary language while slowing down her speech. He thinks that Olga might otherwise have hard time to understand what he says. "Yeah, I'm Russian but I've lived in Finland for seven years. I can speak pretty good Finnish", Olga replies.

Part 19

Sasha, who has followed introductions of Ville and Olga, says to Olga in Russian something that Ville does not understand since he does not speak Russian. Olga looks happily surprised. He answers Sasha right away, also in Russian. However, their Russian-speaking conversation stops abruptly when the doorman opens the door and invites the troupe to step in.

Part 20

The four friends leave their coats in the corner and head straight to the dance floor. They start dancing at the rhythmic pace of the rhytmic techno music.

There is also a quieter side in the nightclub, where one can talk, enjoy one's drinks and eat something little salty, but it will not be seen by Sasha, Villa, Laura and Olga tonight. They go on dancing until the nightclub closes.

Ville and Sasha take turns in entertaining their companions by inventing different imaginative dance moves on the floor. Sasha is even enthusiastic enough throw a volt back when the dance floor is a little less crowded for a moment. It is a failure when his drunken state takes the best out of Sasha's body control.

Part 21

Women respond with their own hypnotic dance moves that make the drunken boys incredibly enthusiastic. Laura and Olga are waving their hips at the same pace in front of Sasha and Ville and sometimes fall into a deep squat.

In spite of his drunken state, Ville remembers to be careful not to dance to Laura. Feeling a little bit frustrated, Ville realizes that he is interested in her much more than Olga, even though Olga is a very beautiful and interesting woman. He feels that Olga senses the situation too. Ville is noticing that Olga is looking more towards Sasha than to him. On the other hand, Ville feels somehow, in a bit of a confusing manner, that Olga might be very interested in him.

Part 22

Ville realizes that he can't really interpret the behavior of a Russian woman. That's why he won't start making any kind of moves on Olga during the evening. He has nothing against the Russians, but he thinks he wants a woman that he can understand.

Nevertheless, all four have an unforgettable fun evening together. Since they are dancing, they are also drinking less. They all are having just a few shots admits dancing.

Part 23

The next morning, Laura is instead Ville in Sasha's apartment. Ville and Olga have taken a taxi because Olga lives in the same direction as Ville. Laura and Sasha have walked through the city with each other holding hands all the way in the autumn night. They slept for a long time, until sunlight amidst passing clouds penetrated into the bedroom and woke them up for preparing breakfast.

Sasha munches his breakfast sandwich and tastes the juice he just squeezed from oranges. He has the newspaper of the morning in his hands. "Well, when one looks at those Baltic countries, they were so much better off to be part of Russia," Sasha says aloud. Laura is in her morning gown at the kitchen pouring coffee into her cup. Her hand stops and an expression of non-belief spreads on her face. "How can you say that?" Laura asks, "are you saying that seriously? " Laura further asks.

Part 24

"How so?" Sasha replies astonished by Laura's tone of voice. "The Baltic countries were really thriving when they were part of Russia, there were great cafes and sandy beaches, people had plenty of food and they were happy. Now, foreign companies have taken the Baltic countries over as their colonies and the Baltic people have to work hard with ridiculously low salaries. In Lithuania, there have been really high suicide rates after they got their so-called independence".

Part 25

Laura cannot believe what she is hearing. She has never read Sofi Oksanen's books about the difficult times in Estonia during the Soviet Union, but she has certainly heard a huge number of stories about the tough inhuman fates that the Estonians were suffering in the decades that are considered the most miserable and distressing in the Baltic history. Sasha's views disturb Laura. She's so surprised she can't say anything. Confused, she pours coffee into her cup and mixes it with a little cream, stirring slowly with a spoon in the cup. For a moment there is silence.

Part 26

Sasha continues his monologue "Finland is a bit different, even though one can see that Finland has sometime been part of Russia. There are a lot of the same words as in Russian and the architecture in Helsinki is similar to St Petersburg. Only that St. Petersburg a bigger and cooler city. Sometime in the past St. Petersburg was the metropolis of Finland, but then there were all those wars. But maybe at some point the Finns will realize to which direction they should look. "After a moment of silence, Sasha continues:" We have to go and watch some year the parade of the Victory Day. Then you'll also see how gorgeously a strong country Russia is. "

Part 27

After that, Sasha opens the morning newspaper. Pages are rustling. "This new tabloid style is much better as I don't the need a large table, to be able to eat while I read the paper," Sasha notes and concentrates on the headlines. His mouth munches the bread again. "Yes, it is quite true," Laura responds, relieved that the difficult topic switched to an easier one.

Laura's delighted voice attracts Sasha. He puts the newspaper aside, rises up and embraces Laura. As he kisses, he whispers to Laura's ear, "My life has been given to me so that I could meet you, lovely Laura." Laura sighs in happiness. She has, of course, heard Sasha say this Pushkin quote before, but somehow it still works. Sasha’s self-confidence and talkativeness have made a strong impression on Laura’s ever since the beginning.

Part 28

Despite the good ending, a concern remains in Laura’s mind over the conversation at the breakfast table. She worries what kind of strange views she will in time discover her Sasha harboring. It is like a lonely cloud drifting on the sky and blocking the summer sun. But no – Laura shakes off her worries. Everything is fine.

Part 29

Ville wakes up in his studio apartment in the outskirts of the city as the sun begins to shine in his eyes. In a flash he remembers the early morning taxi ride. The taxi had stopped in front of Olga’s apartment. Olga had hugged him, given him a kiss and thanked him for the wonderful evening. For a while, Ville had wondered if he should have tried to invite himself over to Olga’s place for afterparty or maybe ask Olga to come to his apartment, but he hadn’t been able to muster sufficient courage. When the taxi rolled off, he had seen Olga disappear into the stairwell of her apartment building.

Part 30

Ville yawns heavily and stretches his limbs pleasurably, gets on his feet and drags himself into the kitchen. After breakfast, he takes a can of beer from the top shelf of the fridge. Without a moment’s hesitation, he opens the can and starts pouring its contents down his throat.

At weekends Ville often does not have any specific program. Rather he spends his time and entertains himself by enjoying beer and wine.

Part 31

Ville can't get Laura off his mind. He recalls the events of the previous night. The blonde beauty's blue eyes, warm smile, velvety soft voice and body shapes revolve around in Ville's mind. "No, I'm not allowed to think about Laura this way, she is my friend's woman," Ville says to himself and goes on to lay on the bed. He reaches for the TV's remote control from the coffee table and clicks open the music channel. On the screen, there are spectacular-looking half-naked young women dancing to techno music.

### Part 32

The vision soothes Ville's mind. He thinks it is best to forget about Laura and go back to the same nightclub, this time alone. He decides to find himself a woman who looks like the girls in the music videos.

However, Ville's plan is going awry. He is in such a bad shape in the evening that the Doorman won't let him inside the nightclub. Ville is then forced to go to a nearby gateway to ease his feeling. The on-site police patrol picks him up from there for a night in a cell.

Ville has been taken to police drunk tank twice before in his life. For the first time, he went a bit too wild in summer festival in inner Finland. The second time he passed out due to excessive alcohol drinking on a park bench on a November weekend night.

### Part 33

The next day, Ville gets out of the drunk tank and immediately calls his friend Sasha. Together, the friends laugh with tiers flowing from their eyes.

This is one of the aspects that makes Ville like Sasha so much. It's not really the first time that Ville has been making a fool out of himself when drunk, but Sasha never condemns his drunken experiences, rather the friends just laugh at them together from the bottom of their hearts.

### Part 34

Ville is not himself worried about his drinking. He believes that his drinking will stop as soon as he finds a woman into his life with whom to settle down. Ville strongly believes that when young, he has to live fully and wildly, seek experiences so that he does not regret his lost youth later. Ville has heard countless stories of middle-aged men who have not been able to live wildly when they were young and have regretted later. The mere thought that he would fall victim to such a fate is very distressing to him.

Part 35

Sasha closes the phone and continues to laugh. “Well, what was all the fun about there?” Laura, listening to the call from the side, asks curiously. Sasha tells her about what happened to Ville the previous day shaking his head and laughing.

“Oh, no, poor Ville,” Laura says compassionately, “and you laugh at it.”

“You should know Ville better so you would understand why we are laughing at what happened,” Sasha explains and continues a bit more serious: “Ville himself laughs at whatever happened to him. It’s like therapy. With me he can share it all, laugh with a friend. And he never gets into any serious trouble. ”Sasha knocks the wooden surface of the table, to make sure nothing more serious will happen to Ville.

Part 36

Laura nods compassiontely. ”Well yeah, a young man sometimes has to try his limits a bit. Great if it has such an ability to laugh at himself. And that he has such a good friend to share with ” Laura says warmly. Then she continues “Guys who always behave correctly are boring. It is only good there is some bad boy in a guy. I did sense in that nightclub that Ville is not a bore. ”

Laura presess herself against Sasha, looking him deeply in his eyes and whispers, “I know you are a bad boy too.”

Sasha slowly lowers his hand to Laura’s waist and kisses her for a long time. Then Sasha lifts Laura up in his arms and carries her to the bedroom.

Part 37

It takes a few days. The first signs of the coming winter are seen in the city. Instead of drizzling rain, big snowflakes start floating down from the sky.

Snowfall is almost hypnotic to watch. Flakes are fascinatingly big ones, and they float down slowly. Occasionally, swirls of light wind catch the snowflakes and fly them up and down for a moment before letting them go. The flakes continue their journey downwards towards the ground, in order to be caught by a new gust of wind just centimeters from the ground. Once on the ground, the flakes will still melt into water, but it is only a matter of time before the fall will turn into a winter.

Part 38

“So wonderful that it is Christmas soon again,” Laura sighs while watching the snowflakes dance in the air. “Sure that is nice, but why are you celebrating it in Finland at the wrong time? And I must say people are celebrating it here a bit comically”, Sasha utters cheerfully.

Laura is startled. Sasha’s harmless jesting gives Laura concerns – what will time reveal of her loved one’s thoughts? “Yeah, yeah I know in Russia you have that frost-man. You have confused the New Year and the Christmas celebrations,” Laura teases.

Part 39

Sasha can’t help feeling hurt. What did Laura mean when she said “You in Russia”? Sasha is Finnish, although his parents moved from Russia to Finland when he was a small baby. Sasha does not like Laura’s voice when she speaks about Russia and Russian culture. There is too much anti-Russianism in Finland. He suspects if his wonderful Laura is one of the many Finns who have suspicions about Russia and Russians?

Sasha is comforted by the thought that Laura has in any case taken her as her partner. They’ve been happily together soon for two months.

Part 40

A silence follows after which Sasha changes the topic of conversation. She asks Laura what her thoughts have been with regards the dinner. Should Sasha go shopping to buy something or what should he do?

Laura says she'd need a little variation today. Sasha feels delighted. He already had a Vietnamese restaurant, a few blocks away, in his mind where portions are super delicious. It is also one of Laura's favorite places. Sasha suggests that he offers Laura a perfect Vietnamese dinner. Laura gets excited – they could try something they haven't tasted before. "Do they serve balut?"

Part 41

In the following days, the temperature drops to ten of degrees below freezing. There is a heavy snowstorm. The snow plows that open the streets push snow to their sides, and cars parked on the roadside are buried in snow. Men and snow-emergency equipment work day and night. Despite that all the snow-plowing capacity is in use, it becomes slower to move about. It seems that the fast approaching Christmas this year will be white, also in the inner city, where in most years there is only slush colored gray-brownish by antifreeze salt that is sprayed on streets.

Snow brings with it more light to the dark time of the year. It stimulates the minds of people moving around on the streets. On the other hand, the cold forces the people outdoors to wrap themselves tightly in their coats. People hurry on their errands as fast as snow and ice on slippery street surfaces allow.

Part 42

One of the persons who finds the streets slippery is Laura, who walks in her hurry towards the university and a lecture she is to attend. She is late because there was a quarrel with Sasha in the morning. Laura is concerned about Sasha's behavior, or rather his personality. Sasha has explained, albeit drunk, that strong emotions are part of him being Russian, that he has a Russian soul that lives strongly and freely.

Part 43

Laura has explained to Sasha that for her, as a Finn, it is important that flames of anger do not constantly flare up. Rather, happiness for her comes via a steady, non-dramatic life.

Part 44

It has been difficult to find a solution to this dissonace. Sasha has often become angry, thinking that Laura is trying to suppress his Russian soul. Even though, Sasha always calms down very quickly, and they have a good moment, the quarrels always leave scars in Laura's soul.

Part 45

Suddenly, there is a really slippery spot underneath Laura's shoe. She loses her balance and falls down painfully on the icy street. Luckily, she does not hit her head, but realizes that she bruised herself. There is pain both in her leg and on her side.

A familiar voice asks with concern: "Hey, Laura are you okay? It was a nasty-looking fall".

Laura turns to see who the speaker was. It is Sasha's friend Ville. Laura blushes slightly. "Ville, is it nice to see you, where did you come from?"

Part 46

Ville explains that he is going to a job interview. He has been unemployed for a long time, things have been slow in the building and development due to economic downturn and also seasonal effect of winter as people tend to redevelop their houses and apartments mainly in the summer. "I am just a working-class guy", Ville sighs.

Part 47

“Working class guy is really cool”, Laura encourages Ville as she to stands up. She is trying her foot is it is good for walking. “I am bothered by Sasha always trying to be some kind of a hero. Maybe it is a part of being Russian. It is just something so stressful”.

Part 48

Laura cannot help to unload what is on her heart, even though a small alarm is tingling at the back of her mind. Is it wise badmouthing Sasha to this good friend?

Ville, however, listens to Laura with compassion. He is wondering if Laura is happy with Sasha. Conflicting feelings rise to Ville’s mind. On the other hand, this is her good friend’s girlfiend, on the other hand, Laura’s dissatisfaction gives him hope.

Part 49

In his eyes, Laura is somehow even more attractive than before. He can’t do anything about it that Laura is attractive to him. And after all, Sasha and Laura have not been dating more than just a bit over two months. What if the relationship does not last long? Then he will be able to ask Laura out. If only…

Ville hides his thoughts carefully from Laura. They’re still exchanging a few words, warmly hugging and wishing each other a good coming Christmas. Both are hurrying to their own ways.

Part 50

Massive white pillars rise to the heights to join the ornate vault arches that support the dome-shaped roof. In the shade of the pillars, there are sculptures that watch with their stony eyes across the main hall of the sanctuary. From the high ceilings hang massive, gold-plated, decorative candlelight lamps. The view is crowned by an altarpiece framed with golden leaf decorations at the front of the hall.

Part 51

The tones of the organ soar. With them, the parishioners in a bit out of sync, but still beautifully, sing Christmas hymns. After the organ stops, the priest, dressed in white long robe, raises a gleaming silver jug and a small silver plate in the light of the chandeliers, and invites the parishioners to come to the offering. Laura leads Sasha into a long queue. “Is it really ok that I am with you here?” Sasha whispers to Laura’s ear. “Of course, Do not be silly now. You have the same God and Savior ”, Laura responds with encouragement. Sasha relaxes.

Part 52

The line is long. Eventually they get to kneel on the altar and receive the offering. “Merry Christmas, my love,” Laura whispers as they return to their seats.

“Is this all? Short delivery, no incense or anything. This church seems to be somewhat poor. But it was nice to be able to sit during the worship”, Sasha whispers.

Part 53

“We have ours this way. In your Russian-Orthodox worship you have your own stuff”, Laura whispers to Sasha’s ear and presses her body against him.

Laura takes a moment to gather her courage and goes on to ask, “Do you feel that this is the kind of service where you could feel comfortable?”

Part 54

Laura is already accustomed to her Sasha having occasionally some quite weird views and thoughts. Laura loves her boyfriend and understands that his parents are Russian. The culture where Sasha has grown, is quite different from Laura’s purely Finnish one.

However, Laura cannot imagine that she would start herself going to Russian-Orthodox worship with Sasha. Those services are so incredibly long and difficult to understand. The worship of icons is also something she does not fully understand. For example, how could a candle lit up on Mary's icon be somehow spiritually meaningful? Laura cannot comprehend that.

Part 55

Laura hopes that for Sasha his Russian-Orthodox background does not mean much. Laura believes that she will make Sasha change his views over time. Perhaps the change started today when Sasha agreed to come with her for the evangelical Lutheran Christmas service. Because of this, Laura becomes very happy when Sasha whispers in her ear: "Yes, I can go here with you if it's important to you."

The most important thing for Sasha is that his Laura is happy.

The end of the Christmas service is coming to a halt, the crowd begins to move towards the exits. Hand in hand, warmly smiling, Laura and Sasha head towards the church exit.

Part 56

The winter sun shines through the window of the trendy city center coffee as brightly as it can in Southern Finland in the last days of December. Light marble floors and glass surfaces of small round tables glitter in the bright spotlight. The white plaster on the walls adds to the effect of this very welcome light phenomenon.

Unlike many other restaurants and cafes, the ceiling of the I is decorated with a light grille with beautiful leaf and floral patterns, instead of bare air conditioning ducts. Metallic chairs painted in black around the small tables create a strong contrast to this light.

Part 57

A shelf runs around the walls of the I with all sorts of historical coffee-related items, old coffee grinders, tin coffee containers, and Southern European espresso pots.

The scent of roasted coffee is seductively floating in the air.

Part 58

Normally a beautiful view on the busy street opens lined on both sides over a hundred years old, decorated houses, opens from the large windows of the I. Right now, however, opposite the coffee shop, I renovation is underway, with ugly scaffolding spoiling the landscape.

Part 59

Laura and Sasha and Laura's good friend Liina are sitting around the furthest window side table. Liina is a cheerful redhead student, about ten centimeters shorter than Laura, dressed in a sleeveless sweater and slightly worn jeans. Liina is Laura's childhood friend who has chosen the University of Art and Design, while Laura went to study law.

Part 60

Hot, steamy appuccinos and delicious croissants are on the table in front of them. The water glasses that come with the Capuccinos have small pieces of lemon that bring a nice, fresh additional flavor to the water. In addition, the table contains brown sugar in a glass container, which they have already put in their appuccinos. Further away in the café, an Arabian-background waitress, dressed in an elegant way, vigorously cleans a table for new customers.

The café is one of the many small, but very positively spirited, venues that have strongly come into fashion in the previous year.

Part 61

Laura feels happy. She and Sasha have spent their Christmas holidays in a calm and relaxed atmosphere. The business of Christmas preparations are behind them. The Christmas party at Laura's childhood home also went fine. The entire family warmly welcomed Sasha and liked his polite, well-behaved being.

This was a great relief for Laura. He was nervous in advance about the way the family might have reacted to his new boyfriend. Laura's boyfriends have changed quite often and some have been outright scoundrels. None of the close relatives have said anything directly, but the reactions have sometimes been bleak.

Laura smiles when she thinks of her former boyfriends. At the same time, she listens to Liina's story about studying in the Art Industry.

Part 62

All of a sudden, a loud rumbling noise coming outside of the café cuts off Laura's thoughts. She turns with other coffee shop customers to look out the window.

On the other side of the street scaffolding had just collapsed on the street. Like a miracle, no one was caught underneath. There are shocked people standing next to the fallen scaffolding. A Mercedes, parked right next to the scaffolding, has been damaged in the accident.

Part 63

"Well, one must say the company that has put up those scaffolding have really done it the Russian style", Laura says, shaking her head in disbelief. Liina laughs sarcastically, but Sasha feels angry in his heart. He always feels bad when Laura talks about something having been done the Russian way. Why does Laura want to talk about Russian culture so very rudely? On the contrary, in Russia, every person in his soul is very careful and conscientious. Does he not understand insulting Sasha when he says so?

Part 64

Sasha hides his hurt feelings from Laura. He remembers well that Laura does not like excessive expression of emotions. Even one spontaneous eruption that, from the point of view of Sasha himself, is just an insignificant flare of temper, can lead to Laura having a bad feeling for the rest of the day.

Part 65

A lonely car drives in the middle of a wintery forest road and rises behind it a cloud of snow. The red Toyota has plenty of speed even though the road surface is frozen. Ville is behind the wheel, he is returning from his parents in the countryside to the city.

Ville is really on a happy mood. He is humming in the pace of the radio “You are a pearl of a hot land, Ramona, Ramona, when I get my arms again, Ramona, Ramona ...”

Part 66

He accurately anticipates every curve of the familiar road, pushes the gas when the road is straight and brakes just before the road bends. Ville’s both hands are firmly on the wheel. He does not need to shift the gears because his parents’ second car has an automatic gearbox that slips smoothly over to the new gear whenever the speed, the slope of the road, and the Ville’s action on gas pedal require.

The fabulous snowy landscapes soothe Villen’s mind. At his parents’ home, Ville has made several walks in the snowy spruce forest surrounding his childhood home. In the dense of the woods, his mind has calmed down and absorbed spiritual energy. In this respect, he is the most typical Finnish person.

Part 67

The snow packed on the spruce branches had been breathtakingly beautiful. On an windless day, one could hear the silence. There was no urban noise pollution within a radius of tens of kilometers. The sound of snow under the shoe and the woodpecker knocking on a tree trunk further in the woods were the only sounds, in addition to the Ville’s own breath and rustle of his clothes. Ville saw the traces of several species of animals in the forest, but wildlife remained at a safe distance from a two-legged traveler and he didn't see any of them.

Part 68

As he is driving, Ville is reflecting how well things are in his life after all. Then he notices a dark figure dart out of the dense spruce forest. The figure runs directly on the road in front of him. Ville can only think "moose" while he instinctively hits the brake pedal all the way to the bottom. The anti-lock braking system vibrates the brake pedal under the sole of his foot. He feels the fast jerking of the studded tires on the icy surface. At the same time, he steers the car towards the side of the road.

Part 69

The moose gallops past Ville's car, the collision is just a few tens of centimeters away. The huge animal disappears across the road to the forest.

Ville pants heavily in the car that has come to full stop. He feels cold sweat in his back and on his forehead. He was very close to losing his life or at least serious injury. The roof of the car he drove has not been made to withstand the collision with the moose at such speeds. It takes awhile before Ville gets his thoughts together enough to continue driving, now just below the speed limit.

Part 70

In his shocked state, a picture emerges in his mind that he can't get rid of: Laura's beautiful, smiling face. "Why don't I get Laura off my mind?" Ville thinks. He shakes his head and another face comes up in his mind, the beautiful Slavic features of the beautiful dark Olga. Ville smiles firmly. He's trying to find Olga in his hands. If Olga gets excited about him, he will be able to forget about Laura.

Part 71

Strong wind blowing from the icy sea to the city penetrates into the bones of even the most well winter-dressed pedestrians. Here and there the wind catches light flakes of snow that have fallen earlier the day. It is a really cold day.

There is a park next to an old church built of red bricks and rising to heights. On a branch of a massive oak, a lonely squirrel looks curiously down and begins an agile descend along the tree trunk. Something attracts him. The squirrel stops occasionally on his way to sniff. There is something something good below.

Although there are hardly any enemies of squirrels in the city, sometimes little boys have tried to catch the squirrel. Dogs kept free in the park may also be excited to chase a small furry creature. Those times have consolidated in the squirrel's memory. Therefore, he is wary of his surroundings on his way down the from the oak.

Part 72

Below, on a park bench is sitting a dark-haired and brown-eyed woman clad in a thick winter fur. The woman, passing the oak, has noticed the squirrel in the tree. In her pocket, there have been nuts she has taken in her hand. Now, she is handing the nuts towards the squirrel and waiting patiently.

The squirrel is approaching, sometimes stops with suspicion, but the nuts attract him too much. The woman is patient to remain stationary. At last, her patience is rewarded: the squirrel reaches for her hand, grabs a nut, and then quickly flees about one and a half meters away to eat the nut. At the same time, he eyes the woman with his squirrel eyes. "Oh, he is so cute", says a quiet male voice behind the woman. The woman is slightly startled, raises and turns to look back at the man who has appeared behind her. "Hey, I remember you," she says in Russian. The man, who is Sasha, replies in Russian: "Yes, I remember you, you are Olga, from that nightclub queue, Laura's friend from the university,  us dancing that night together."

Part 73

Sasha is pleasantly surprised to come across a familiar person. There is also a warm smile on Olga's face. He hugs Sasha. The memory of the handsome Russian youth has been left permanently into Olga's mind after those hours spent on the nightclub dance floor. She has secretly hoped to meet Sasha again. Olga therefore suggests that they go to some warm coffee shop to talk. She burns in desire to know where from Russia Sasha’s parents are from, and to get to know him better.

Part 74

To Olga's great pleasure Sasha agrees right away and promises to offer Olga coffee and some pastry. Olga throws the rest of the nuts in the snow in front of the squirrel. They then look together, as the squirrel comes closer, stops and grabs the nuts in his small little paws. Then they look at each other, and head for a café in a Jugend-style house on the edge of the park.

Part 75

Ville trembles amidst the cold snowstorm. He has gone for a walk and traveled much further than he originally intended. The cold has begun biting his toes and fingers, not to mention his nose.

Ville has a problem. He had been thinking of calling Olga and asking her out. For that, he had found out Olga's surname and her phone number. But Ville is nervous. A number of questions revolve in his mind. Would Olga even remember him? Would there ever be chance of success? Ville remembers all too well how everything went wrong in his short relationship with Svetlana. He simply failed to understand Svetlana's thinking. Ville does not know whether the problems were due to the Svetlana’s personality or her being Russian. Would Olga be difficult to understand in the same way? Ville and Svetlana, after only a couple of weeks of fiery love, decided together that they were not meant for each other, despite the fact that they still had strong feelings toward each other.

Part 76

When thinking about these issues, Ville picks up his smartphone from his pocket and starts dialing Olga's number. Then his fingers stop, and he looks up at the night sky. Snowflakes float down on his face. Ville closes his eyes, presses down his head and shakes it slowly. Then he wipes the screen of his phone clean from the snow and slides the phone back into the pocket. "What should I do?" He asks himself quietly and then continues the slow walk. Snow crunches under his shoes. The snowstorm will soon cover his footprints.

Part 77

Spring is coming to the city center. After the long seasonal dark period, the sun is glimmering behind the clouds, the spring rain has stopped, and the happy twitter of birds fills the fresh air.

Late in the afternoon, after the lectures are finished, Laura rushes to her beloved Sasha's one-bedroom apartment, where Sasha is already waiting for her. Sasha has bought tasty roast meat, which Laura has skillfully marinated and put on the baking pan with carrots, onions and chopped potatoes. Laura enjoys cooking and is skillful, which really suits Sasha.

Part 78

After putting the baking ingredients into a hot oven, Laura starts making salad and salad dressing. Laura's phone starts ringing in the middle of this activity. Laura quickly flushes her hands, dries them with the kitchen cloth and answers the phone.

The caller is her friend Liina, who has heartache. There has been a breakthrough between Lina and her boyfriend who is studying graphic design. The boyfriend has found another woman and left Lina. Liina suspects that this other woman is one of their common student friends.

Part 79

"Oh no, that is terrible," Laura exclaims and comforts Liina to her best. Laura knows that empathic listening is the best help in these situations. She puts her hand on the phone for a moment and whispers to Sasha that Liina's boyfriend has left her. Sasha communicates with his expressions to Laura that she is sorry for Liina.

Both know that the first days are the worst. After that, recovery begins. In the best case, there is soon a new affection and a new opportunity to sweep away the bitter disappointment of a losing relationship.

Part 80

Suddenly Laura is alerted by the scent of smoke. She has been lost in her phonecall with Liina and has not noticed the passage of time. Baking ingredients in the oven have begun to burn. "Oh, no, now I've not totally messed this food the real Russian way", Laura pains. She takes the roast from the oven and at the same time apologizes to Liina. "We have to quit now," Laura says, closes the phone and glances at Sasha as with feelings of quilt. Laura sees Sasha being upset. "I'll try to remove the burnt surface. Maybe it can still be eaten. "

Part 81

However, Sasha is not annoyed by the spoiled roast. He can no longer restrain himself but raises his voice: "You always say how this and that has been done the Russian way. It is a really upsetting, you might think a little about the feelings of the other, even if you think that my home culture is a bad one ".

Laura looks at Sasha with astonishment, "I don't mean anything with it, it is just an expression".

Sasha is not calmed down by this, but with a few hurried irritated steps he is in the hallway, puts his shoes on his feet, jerks his coat from the coathanger so that Laurank's coat flies on the floor and leaves the apartment. The door slams shut behind him.

Part 82

Laura feels really bad when she realizes she has offended Sasha. At the same time, she is shocked by how childish and, on the other hand, even intimidating temperament, Sasha possesses. Laura is in despair. Is Sasha then the right one for her? Their temperaments are so different. Would she ever come to be in terms with Sasha's fiery nature? Could Sasha change? Would love between them last? Laura's limbs suddenly feel heavy like lead, and she sits down on a stool in the hallway. Tears swell into her eyes.

Part 83

Laura's phone starts to ring again. Laura gets up and retrieves the phone from the kitchen, believing Liina is calling her again. Laura's heart races when she sees that the caller is not Liina, but Ville. Laura takes a couple of deep breaths to get her voice steady and then respond “Laura”. Already when she says that, she realizes that Ville would hear the quiver of her voice.

“Well hey it is Ville here, I tried to call Sasha but he doesn't answer. Is he there somewhere? ”Ville asks. He pauses and then continues, worried: "Your voice sounds like you’ve been crying ... is all right?"

Part 84

Laura is no longer able to hold her feelings inside. She tells, sobbing, about their argument with Sasha. Telling Wool makes it somehow feel easier to her. He listens understandingly. Laura is reminded of how balanced and calm a personality Ville is, how he has taken things calmly in all circumstances without ever getting angry. Ville also knows how to laugh off his drunken mishaps from his mind.

Ville asks if Laura would want to go to a nearby cafe to talk about it. Without thinking for a moment, Laura agrees. She is surprised to see how much she wants to talk to him, in particular, about her situation.

Something transformative has taken place in Laura's mind. Ville is attracting her now. Laura suddenly realizes, in a flash, that her life has changed its direction.

Part 85

Laura knows that Sasha is quick to cool down, but she doesn't want to be there anymore when Sasha comes back. They agreed with Ville to meet in a familiar restaurant a couple of blocks in quarter of an hour.

Laura glances at the mirror and quickly fixes her makeup so that her face doesn't look like she had been crying. She lifts her red coat off the floor and pulls it over herself, chooses high-heeled leather boots from her shoe rack and exits to the staircase.

Part 86

Evening has already fallen. It is misty, gray and cool. A drizzle has slowly made the asphalt and rocks of the streets moist. Lights from windows of street level shops and the dwellings above them cut through the darkness of the evening. Water rushing to the streets from downspouts form puddles on the lanes. The engines of rarely passing cars are buzzing, puddles are splashing as they run over, and the last studded winter tires of the spring make their noises. There's a fresh scent of early spring in the air.

Part 87

Laura's high-heeled boots are echoing on stones of the pavement as she slips under the entrance of the cafe in the street corner. Laura closes her umbrella and shakes off the excessive water on the street before stepping into the coffee shop.

Part 88

Ville sits at a wooden round table in a back corner of the cafe and tastes heated wine. He had surprised himself as he proposed a meeting to Laura. On one hand, Ville has a bad conscience for approaching Laura, the girlfriend of his friend. On the other hand, he thinks that if there are some mutual feelings between him and Laura, therefore wouldn't it be right for them to end up together. No man can help himself when in love. "A man in love is like a tree that has fallen into a flooding stream," Ville thinks. "You simply have to go where the stream takes you."

Part 89

If Laura really cares about Sasha, Ville doesn't have any chance of getting Laura to himself. However, Ville feels he has given this a try. He is ready to end his friendship with Sasha if it is the price to pay for love.

Ville feels his heart jumping when he sees Laura sneaking in via the door of the cafe. Laura looks around. Then the cute eyes flash and the warm smile lights up when Laura notices Ville in the corner table.

Part 90

Laura has a short skirt and a thin blouse with ample cleavage, but Ville sees only Laura's radiant blue eyes and her enthusiastically smiling face. Suddenly Ville realizes that Laura is no longer sad. A happy smile rises to Ville's face and he is not able to anything other out of his mouth than: ”Hi! So nice you came. ” Laura is sitting down and is saying anything. They just look at each other deep in the eyes. Then their faces get closer to each other. Lips press against each other and open for a long, lingering French kiss.

Part 91

April has come. Morning sun shines on clean white linen at Ville’s apartment. Laura and Ville rest on the bed, intertwined, slowly awakening in to the late spring morning. Laura smiles, plays with Ville’s short hair and rises to sit on the edge of the bed.

Part 92

"I'll make breakfast for you, rest for a while." This makes Ville smile too. “You are such an awesome woman. Prepare a herring sandwich for me, and strong coffee too. That will help me overcome my hangover”. “You should not drink so much” Laura scolds Ville with a motherly smile. “You just love me for being a scoundrel like this”, Ville laughs. Laura grabs a pillow from the bed and, still smiling, hits Ville in the face. “Okay okay”, Ville laughs, “I’ll stop drinking altogether”.

Part 93

He does not believe even himself in what he says. But Laura knows better. She goes to kitchen to prepare the breakfast. First, she prepares the sandwich that Ville wished for. Then she takes some cereals and natural yoghourt. Laura finishes off the preparations by pouring into two glasses ice-cold orange juice from the fridge.

Next, Laura puts on the coffee brewer. The day before she has bought some ecological coffee that Liina had warmly recommended. Now Laura waits eagerly to get to taste that.

Part 94

Laura is happy to have found Ville into her life. It has been four weeks since their kiss in the cafe. After that, Laura no longer wanted to return to Sasha's apartment. They went straight over to Ville’s from the cafe. Necessary arrangements were taken care of later with Sasha.

Laura stops to remember that first evening and the night that followed. Ville has a bad boy side to him that has a strong appeal to Laura. At the same time, however, she clearly sees that, as Ville is madly in love with her, she would be able to transform Ville to be decent enough guy to be a good husband and father of their children.

Part 95

Suddenly, Laura realizes that the aroma of coffee does not smell good, but rather she feels it is repulsive. Laura is startled as emotions rush in. Is this why she has been in vain to wait for her menstruation to begin?

Laura grabs her handbag and slips into the bathroom. After a while, she returns to the kitchen with a thoughtful look. She takes a coffee pot and pours it, but only in Ville’s cup. She leaves her own empty. As

Laura is engaged in these preparations, Ville creeps behind Laura with an intention to surprise her. However, Ville immediately sees that something has happened.

"Is everything all right?" Ville asks. Laura looks Ville straight in his eyes and says, “I'm pregnant. You will be a father ”.

Part 96

For a moment Ville fails to understand what Laura says. Father? Him?

"Is this really true?", Ville finally manages. "Yes, a real baby is on the way," Laura replies, trying to interpret Ville's reactions at the same time. Laura feels fear in her stomach. In her mind, she worries if this relationship moved too quickly to this stage.

However, Laura's fear is unjustified. In that moment, Ville realizes that he has now got Laura really to himself. When they have will have a child, no one could come in between them as long as he only takes good care of Laura and the child and their mutual love. Ville starts to smile radiating happiness.

"This is unreal," he says and hugs Laura. “Now we have to open a bottle of champagne to celebrate” At the same instant, Ville realizes that Laura can no longer drink alcohol. Ville and Laura burst into happy laughter together.

Part 97

Saturday in May is sunny and warm. Sasha and Olga are sitting on a park bench, Olga is in Sasha's lap. She has wrapped her hands around Sasha's neck. Sasha caresses Olga's long dark hair. They look at each other deep in the eyes so close that their noses gently touch each other. The world about them is pushed to the background, words are not needed.

Part 98

Intermittently, the tips of their tongues gently meet as they kiss. Then they immerse themselves again to explore each other's eyes, the mirrors of soul.

By them walks an old woman who has been pressed into a hunch by the countless years. She stops to watch the sight. A happy smile spreads on her face. She is muttering quietly to herself, “This is what the spring does… nice to see when young people are in love.” Then she sighs and happily continues her journey.

Part 99

Sasha has bought a ring earlier in the day. Now he encourages his mind and presents the most important question of his life. Olga looks at him for a moment and answers in Russian: “Yes Sasha, my loved one. Yes…"

Their story began on that snowy evening when Sasha found the lovely Olga in the park feeding the squirrel. They had been talking in Russian until the closing time of the coffee shop. Sasha and Olga were then both realizing that they were meant for each other, they were soul mates. Still, they both tried to fight those feelings because Sasha was with Laura.

Part 100

In the evening that Laura accidentally burned the roast in the oven and irritated Sasha by talking about having done it the Russian way had resolved the situation. Sasha returned home, but Laura wasn't there. Eventually, there was a text message from Laura, in which Laura said that she could no longer continue the relationship. Sasha had instantly called Olga and Olga came to Sasha right away.

Since that evening, they have spent as much time together as possible. They are connected by many things big and small, love of Russian culture, language, food, similar temperament, common values. It is like they had been really created for each other. Now they are looking forward to happy summer wedding and are planning ahead to family life. Sasha's happiness is crowned by the idea that their wedding will be held with the warmest possible Orthodox Christian manner.

Part 101

Yes, and about that Ville… Well, the Russian soul is great. One should not fixate, but rather forgive and live on. For what else does a person have besides the wonders of his life?

The spring sun feels wonderfully warm to Sasha. He embraces Olga.

That is where they now are, sitting on the park bench, in love.

Could life be more wonderful?
